# Live-cell single RNA imaging reveals bursts of translational frameshifting

**DOI:** 10.1101/478040

**Authors:** Kenneth Lyon, Luis U. Aguilera, Tatsuya Morisaki, Brian Munsky, Timothy J. Stasevich

**Affiliations:** Department of Biochemistry and Molecular Biology, Institute of Genome Architecture and Function, Colorado State University, Fort Collins, CO, 80523, USA; Department of Chemical and Biological Engineering and School of Biomedical Engineering, Colorado State University, Fort Collins, CO, 80523, USA; Cell Biology Unit, Institute of Innovative Research, Tokyo Institute of Technology, Nagatsuta-cho 4259, Midori-ku, Yokohama, Kanagawa, 226-8503, Japan

## Abstract

Ribosomal frameshifting during the translation of RNA is implicated in both human disease and viral infection. While previous work has uncovered many mechanistic details about single RNA frameshifting kinetics *in vitro*, very little is known about how single RNA frameshift in living systems. To confront this problem, we have developed technology to quantify live-cell single RNA translation dynamics in frameshifted open reading frames. Applying this technology to RNA encoding the HIV-1 frameshift sequence reveals a small subset (~8%) of the translating pool robustly frameshift in living cells. Frameshifting RNA are preferentially in multi-RNA “translation factories,” are translated at about the same rate as non-frameshifting RNA (~2 aa/sec), and can continuously frameshift for more than four rounds of translation. Fits to a bursty model of frameshifting constrain frameshifting kinetic rates and demonstrate how ribosomal traffic jams contribute to the persistence of the frameshifting state. These data provide novel insight into retroviral frameshifting and could lead to new strategies to perturb the process in living cells.

## INTRODUCTION

Frameshifting is a fundamental biological process in which a ribosome translating an RNA slips by +/-1 nucleotides, resulting in the translation of an entirely different peptide sequence from that point forward. While frameshifting is generally detrimental to protein fidelity (Belew et al., 2014; Choi et al., 2015), the process effectively creates two distinct proteins from a single RNA (Clark et al., 2007; Meydan et al., 2017; Yordanova et al., 2015). Viruses exploit this aspect of frameshifting to minimize their genomes and to successfully replicate in host cells (Brierley and Dos Ramos, 2006; Brierley et al., 1989; Caliskan et al., 2015; Cardno et al., 2015; Mouzakis et al., 2013). A prototypical example is HIV, which utilizes frameshifting to translate the gag-pol proteins from a single viral RNA (Guerrero et al., 2015).

Although frameshifting has been extensively studied *in vitro* and in bulk assays (Caliskan et al., 2014; Charbonneau et al., 2012; Chen et al., 2014; Lopinski et al., 2000; Mouzakis et al., 2013; Ritchie et al., 2017), the process has never been observed at the single molecule level in living cells. This leaves many basic questions about frameshifting unresolved. In particular, it is not clear how heterogenous frameshifting is from one RNA to another, nor is it clear if single RNA continuously frameshift in a constitutive fashion or if instead they frameshift in definable bursts, as has been observed for transcription (Lionnet and Singer, 2012) and 0-frame translation (Wu et al., 2016). Finally, the localization of frameshifting has never been investigated, so it is not clear if frameshifting occurs all throughout the cell or is instead preferentially localized to specific sub-cellular regions. In the case of HIV-1 gag-pol, for example, previous assays have shown that 5-10% of translated protein product is frameshifted (Brierley and Dos Ramos, 2006; Dulude et al., 2002; Grentzmann et al., 1998; Mouzakis et al., 2013). This is thought to occur when ribosomes translate through a specialized frameshift sequence (FSS) containing a stem loop structure that slows down incoming ribosomes and causes them to slip back one nucleotide on a slippery sequence preceding the stem loop. Whether or not this occurs constitutively and with equal probability on all HIV-1 RNA or if instead it occurs on a specialized subset that are in the right place, at the right time, and with the right factors remains to be determined.

To directly address these sorts of questions, we have developed technology to visualize and quantify single RNA frameshifting dynamics in living cells. Using multi-frame repeat epitopes, complementary high-affinity fluorescent probes that selectively bind the epitopes, and multicolor single-molecule microscopy, we are able to simultaneously monitor the translation of single RNA into two unique nascent polypeptide chains encoded in shifted open reading frames. Application to the HIV-1 FSS uncovers unexpected heterogeneity in the production of frameshifted product and implicates a novel bursty frameshifting mechanism. Besides frameshifting, our technology can now be used to examine other translational regulatory dynamics, including upstream open reading frame selection, non-canonical initiation, and ribosomal shunting. We anticipate multi-frame nascent chain tracking will be a powerful new tool to dissect complex translational regulatory dynamics in living cells and organisms.

## RESULTS

### A multi-frame tag to monitor single RNA translation in two reading frames simultaneously

We created a multi-frame (MF) tag to monitor, in living cells, the translation of single RNAs with overlapping open reading frames (ORFs). The tag builds off earlier technology to visualize translation using repeat FLAG or SunTag epitopes labeled by fluorescent Fab or scFv, respectively (Lyon and Stasevich, 2017; Morisaki and Stasevich, 2018). In the MF tag, FLAG epitopes in the 0 frame are separated from one another by SunTag epitopes in the −1 frame. With this arrangement, single RNAs with ribosomes translating the 0 frame will produce FLAG epitopes labeled by Fab, while those with ribosomes translating the −1 frame will produce SunTag epitopes labeled by scFv. Thus, depending on the chosen frame(s), polysomes will appear all green (all ribosomes translating the 0 frame), all blue (all ribosomes translating the −1 frame), or some combination of the two (Fig. 1A).

**Figure 1.**
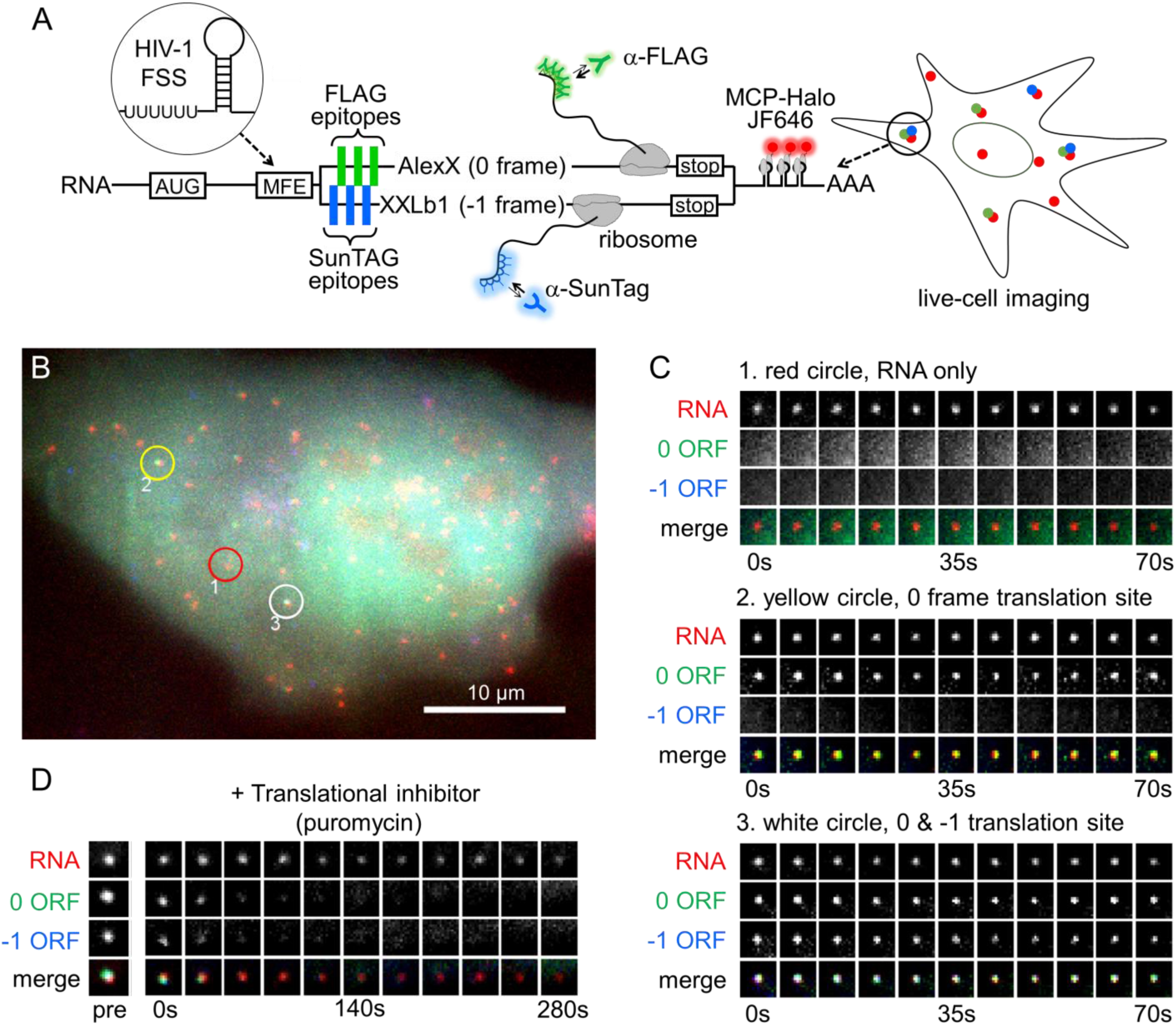
A multi-frame tag to image single RNA frameshifting dynamics in living cells. (A) The multi-frame (MF) tag contains 12 repeated FLAG epitopes in the 0 frame interspaced between 12 repeated SunTag v4 epitopes in the –1 frame. Depending on which frame is translated, nascent epitopes are labeled by fluorescent anti-FLAG antibody fragments (Cy3-Fab, green ‘Y’) or anti-SunTag single chain variable fragments (scFv-GFP, blue ‘Y’). Following the repeat epitopes is exon 1 of the GNAS locus, in which the peripheral membrane proteins AlexX (689 aa) and XXLb1 (690 aa) were placed in the 0 and −1 frames, respectively. Preceding the multi-frame tag is a multi-frame element (MFE). In this study, the HIV-1 frameshift sequence was used as the MFE. To facilitate single-RNA tracking, a 24X MS2 stem-loop tag was also placed in the 3’ UTR. This tag is labeled by MCP-HaloTag (with JF646-HaloTag ligand, red). (B) A representative cell ~10 hours after transient transfection with the multi-frame tag depicted in A. The red circle (labeled ‘1’) highlights a non-translating RNA, the yellow circle (labeled ‘2’) highlights a 0- frame translation site (TS), and the white circle (labeled ‘3’) highlights a 0 & −1 TS. Scale bar, 10 µm. (C) Montages showing the temporal evolution of the RNA spots circled in B. (D) A representative montage showing the loss of signal from the 0 and −1 open reading frames upon addition of the translational inhibitor puromycin (100 µg/mL).

To ensure both frames of the MF tag encode functional proteins and have similar ribosomal occupancy, we inserted the first exon of the human GNAS1 locus downstream of the FLAG and SunTag epitopes. The GNAS1 locus contains two overlapping ORFs of roughly equivalent lengths that encode peripheral membrane proteins in the 0 and −1 frames: XXLb1 and AlexX (Abramowitz et al., 2004; Aydin et al., 2009) (Fig. 1A). In combination with the epitopes, this arrangement has several advantages: 1) epitope densities are maximal since −1 frame epitopes act as linkers for the 0 frame and vice versa; 2) signals are digital, so frameshifted and non-frameshifted species are marked by two distinct probes/colors, and 3) epitopes are placed in nearly equivalent positions, so signals appear at roughly the same time and with similar amplification when translated with similar kinetics.

As a first application of the MF tag, we focused on −1 programmed ribosomal frameshifting caused by the HIV-1 frameshift sequence (FSS). We inserted the FSS upstream of our MF tag and transiently transfected the resulting construct into U-2 OS cells. The FSS contains a slippery poly-U stretch nine nucleotides upstream of a stem loop. Two to ten hours post transfection, we observed cells with tens or hundreds of individual RNA diffusing throughout the nucleus and cytoplasm (Fig. 1B). Nascent Chain Tracking (NCT) (Morisaki et al., 2016) of the RNA revealed a high degree of RNA-to-RNA heterogeneity, with a subset of RNA labeled by Fab only – indicative of FLAG epitopes from translation in the 0 frame – and a smaller subset labeled by both Fab and scFv – indicative of both FLAG and SunTag epitopes from canonical and frameshifted translation in the 0 and −1 frames, respectively (Fig. 1B,C and Movie S1).

To confirm these RNA were active translation sites, or polysomes, we performed two experiments. First, we re-imaged cells 12-24 hours after transfection. At these later time points, Fab and scFv began to accumulate in the cell membrane (Fig. S1 and Movie S2, left panels), as would be expected if they labeled mature and functional XXLb1 and AlexX proteins (Aydin et al., 2009). In the control cells without the FSS, no significant accumulation was seen from the −1 frame (Fig. S1 and Movie S2, middle panels), despite this frame encoding a functional protein, as demonstrated by shifting the sequence by one nucleotide into the 0 frame (Fig. S1 and Movie S2, right panels). Second, we treated cells with the translational inhibitor puromycin. Just minutes after treatment, we observed a dramatic decrease in the number of Fab- and/or scFv-labeled RNA, consistent with the premature release of nascent chains (Fig. 1D and Movie S3). Together, these data provide strong evidence that we are able to detect single RNA frameshifting dynamics with the MF tag.

### Using the multi-frame tag to quantify HIV-1 frameshifting efficiency

The HIV-1 FSS structure has been previously shown to produce frameshifted protein with an efficiency of 5-10% based on the dual luciferase assay and similar bulk assays (Brierley and Dos Ramos, 2006; Dulude et al., 2002; Grentzmann et al., 1998; Mouzakis et al., 2013). However, it remains unclear how this percentage is established. One possibility is that all RNA behave more or less the same and their ribosomes frameshift with 5-10% probability. On the opposite end of the spectrum, it is possible that RNA display a high degree of heterogeneity, so that just 5-10% of RNA have ribosomes that frameshift with nearly 100% probability. A third possibility is that frameshifting is common on all RNA, but frameshifted proteins are less stable and degraded faster than non-frameshifted proteins.

As a first step to quantify frameshifting dynamics at the single-molecule level, we tracked thousands of individual RNA approximately two to ten hours post transfection with and without the FSS sequence present (±FSS; Fig. 2A-C). In the +FSS cells, we found 92 ± 1.3% of translation sites were translating the canonical 0 frame alone, while 6.2 ± 1.1% were translating both the 0 and −1 frames. Only rarely did we observe translation sites translating just the −1 frame (1.6 ± 0.5%, Fig. S2 and Movie S4). To ensure these results were not influenced by the multi-frame tag, we reversed the FLAG and SunTag epitopes in the tag and repeated experiments. This gave nearly the same percentages, confirming the tag order and/or epitope positioning did not bias measurements (Fig. S3 and Movie S5). We then repeated experiments in cells transfected with the control -FSS construct. In this case, we observed virtually no frameshifting sites (0.9 ± 0.7%) (Fig. 2C and Movie S6). Taken together, these data suggest the FSS alone causes ~8% of translation sites to frameshift.

**Figure 2.**
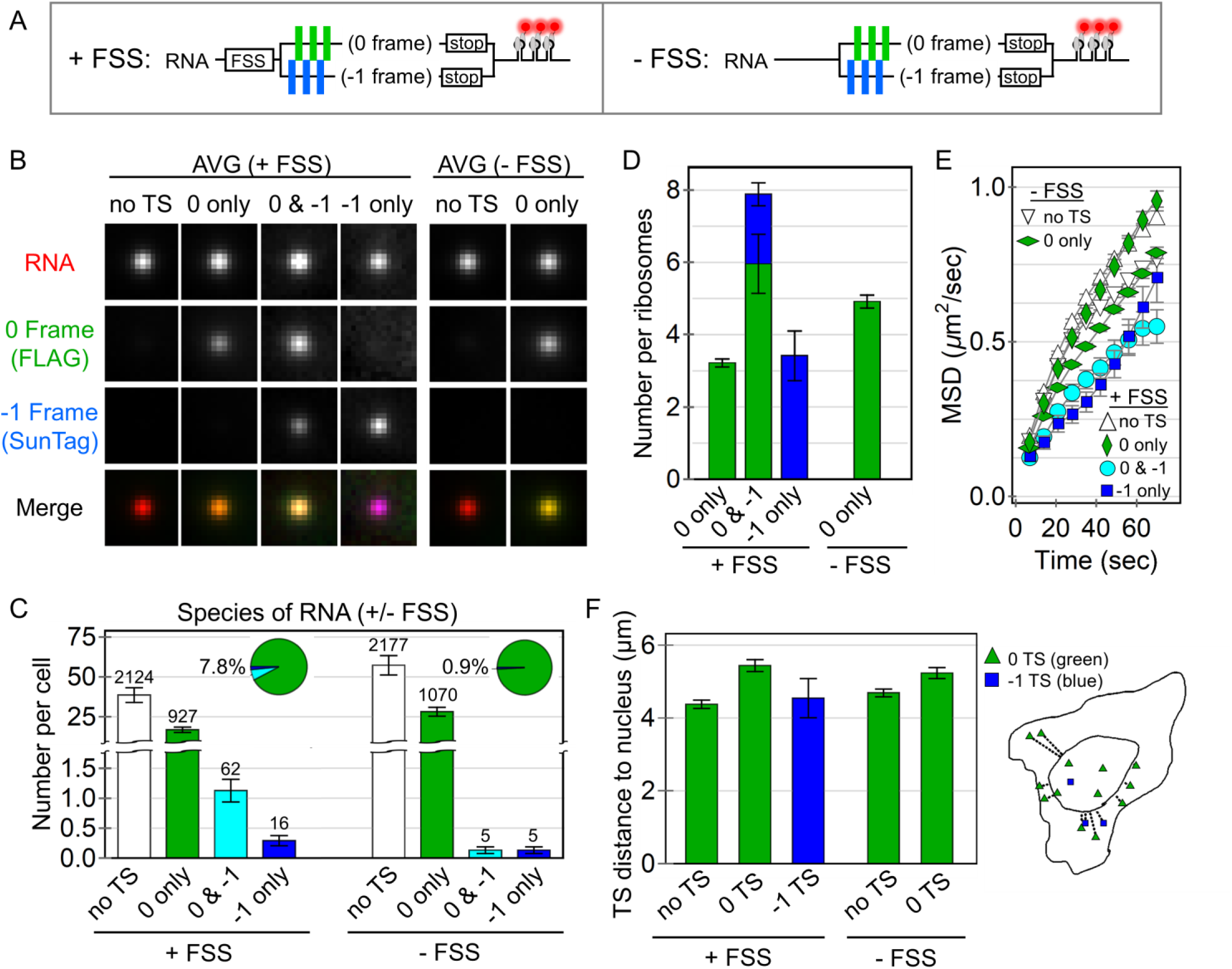
Quantification of HIV-1 stimulated frameshifting. (A) A schematic of the multi-frame (+FSS) and control (−FSS) reporters. (B) Average image trims of all non-translating RNA sites (no TS), 0 frame translation sites (TS), 0 & − 1 TS, and −1 TS, with their respective merges. (C) The number of RNA detected per cell transfected with either the +FSS reporter (60 cells, 3129 total RNA) or the control -FSS reporter (49 cells, 3257 total RNA). The pie charts show the percentage of translating species per transfected cell. The number of detected RNA for each species is shown above each bar. Error bars represent S.E.M. among cells. (D) The number of detected ribosomes per translation site for the +FSS and –FSS reporters. Error bars represent S.E.M. among RNA. (E) The mean squared displacement (MSD) of tracked RNA species as a function of time. Error bars represent S.E.M. among RNA. (F) The average distance (µm) of detected translation sites from the nucleus. An outline of a representative cell on the right shows all detected RNA within the cell and their measured distance from the nuclear border (inner curve).

To further characterize the efficiency of frameshifting, we quantified the precise number of frameshifted versus non-frameshifted ribosomes per translation site. To do this, we re-imaged the MF tag at high laser powers so we could simultaneously visualize single mature XXLb1 or AlexX proteins in comparison to their nascent chains within translation sites (Fig. S4). After renormalizing the fluorescence, we found frameshifting sites with translation in both the 0 and −1 frames had 7.8 ± 0.7 ribosomes total, 1.9 ± 0.3 of which were frameshifted, while 0-frame only translation sites had 3.2 ± 0.1 ribosomes and −1-frame only translation sites had 3.4 ± 1.4 frameshifted ribosomes (Fig. 2B,D). Thus, although just ~8% of translation sites contained frameshifted ribosomes, within these subset of sites, anywhere between 30 to 100% of ribosomes were frameshifted. These data support a heterogenous RNA model in which frameshifting occurs on a small subset of RNA with high probability.

### Frameshifting occurs preferentially in multi-RNA “translation factories”

Given the heterogeneity of observed frameshifting, we hypothesized that the frameshifting state could be stimulated by a specific sub-cellular environment. To test this hypothesis, we performed a statistical analysis of all tracks to see if any biophysical parameters correlated with frameshifting. This revealed frameshifting sites diffuse more slowly than other translation sites or RNA (Fig. 2E) and were also slightly more likely to be near the nuclear periphery (Fig. 2F). These results are consistent with a slight preference for frameshifting in the perinuclear endoplasmic reticulum, where RNA are less mobile and more efficiently translated (Voigt et al., 2017; Wang et al., 2016).

Of the parameters we quantified, one of the strongest correlates of frameshifting was RNA intensity. Specifically, sites translating the −1 frame had an average RNA intensity that was nearly 30% brighter than RNA only spots (Fig. 2B and S5A). In addition, these brighter frameshifting sites were translation-dependent, dissociating upon puromycin treatment (Movie S7). We wondered if this might be an artifact due to the aggregation of probes. Self-aggregation of Fab was unlikely as this would cause 0-frame translation sites to also be brighter, which was not the case (Fig. 2B). Similarly, it was unlikely to be due to self-aggregation of scFv, as this would cause 0-frame translation sites to be brighter in the reverse MF tag, which was also not the case (Fig. S5B). This left the possibility that Fab aggregate with scFv. To rule this out, we reimaged the MF tag without Fab. As frameshifting translation sites remained brighter than other sites (Fig. S5C), we conclude the brighter RNA signal is not a tagging artifact, but instead represents a propensity for frameshifting sites to associate with other translating RNA in higher-order complexes reminiscent of “translation factories” (Morisaki et al., 2016; Pichon et al., 2016).

We did not observe factories in cells transfected with the control -FSS construct (Fig. S5D). We therefore reasoned the FSS itself might facilitate frameshifting in factories. To test this model, we co-transfected a short oligo RNA containing the FSS alone into cells expressing the +FSS MF tag. This led to a significant increase in the fraction of frameshifting sites translating just the −1 frame, from 1.6 to 5.6% when 1 μg oligo was added, and up to 9.5% when 4 μg oligo was added (Fig. 3A,B). The significant increase in the −1 only sites suggests RNA can exist in a state in which all or nearly all ribosomes are frameshifted. Moreover, frameshifting continued to occur preferentially within factories, having RNA signals from 30 to 50% brighter than RNA-only spots (Fig. 3C), even though the overall fraction of RNA within factories remained unchanged (Fig. S6). Importantly, in control cells expressing the -FSS construct, we did not see an increase in frameshifting upon oligo RNA co-transfection. Collectively, these data provide further support for a model in which the HIV-1 FSS itself facilitates frameshifting in factories.

**Figure 3.**
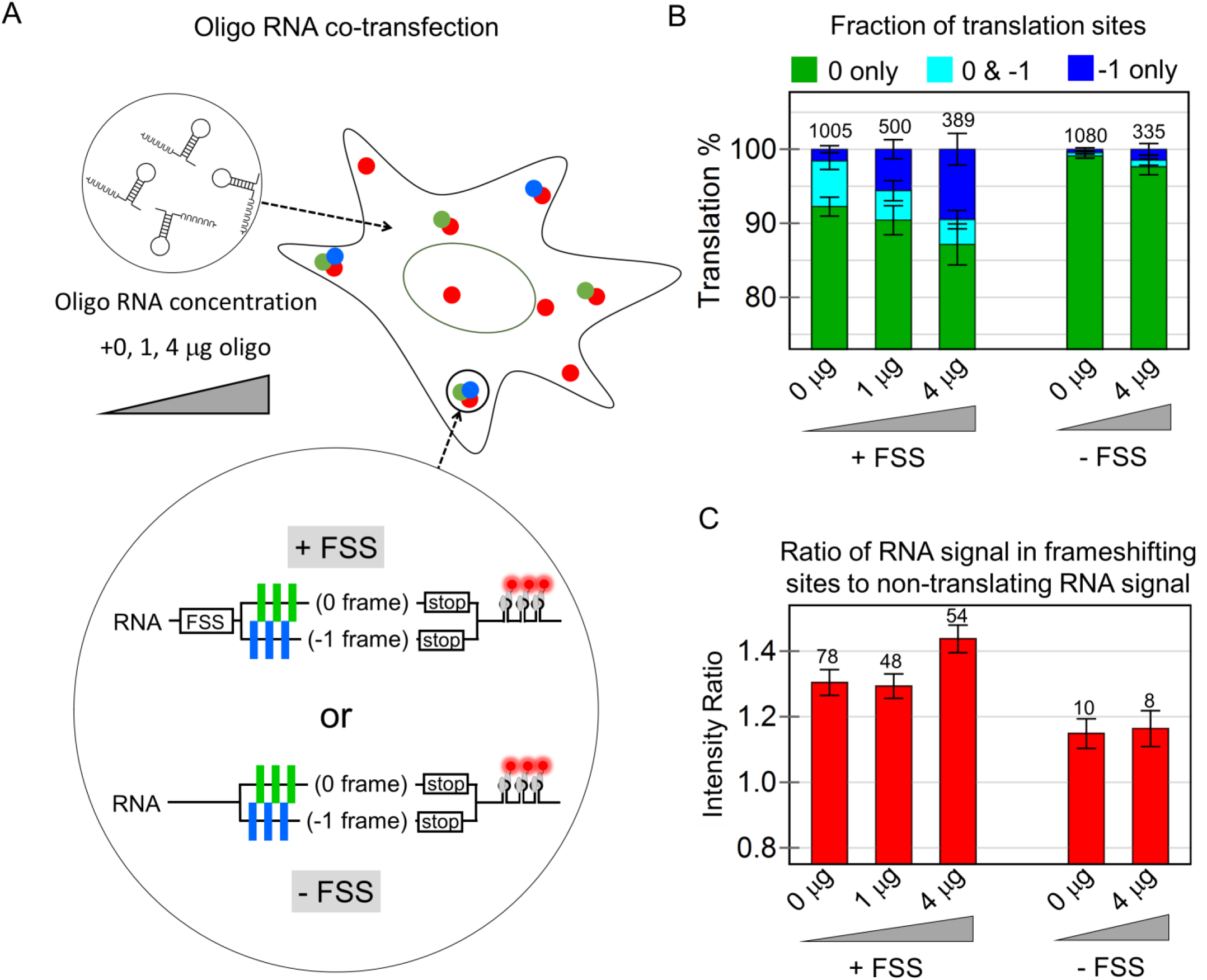
The frameshift sequence stimulates frameshifting in multi-RNA translation factories. (A) Cells were co-transfected with increasing concentrations of short oligo RNA encoding just the frameshift sequence, together with either the multi-frame reporter (+FSS) or control reporter (−FSS). (B) The percentage of translation sites translating just the 0 frame (green), the 0 and −1 frame (cyan), or just the −1 frame (dark blue) at all oligo concentrations for the +FSS and –FSS reporters. (C) The ratio of the intensity of RNA signals within frameshifting sites to non-frameshifting RNA at all oligo concentrations for the +FSS and -FSS reporters. In all bar graphs, the total number of each type of RNA species is shown above each bar and the error represents S.E.M. between cells.

### Translational output of frameshifted ribosomes

The ~8% of frameshifted translation sites we observed is consistent with previous measurements of 5-10% frameshifted protein product (Brierley and Dos Ramos, 2006; Dulude et al., 2002; Grentzmann et al., 1998; Mouzakis et al., 2013). All else equal, this implies that frameshifting alone can explain the steady-state levels of frameshifted protein, without the need for other regulatory mechanisms, such as protein degradation. To test this hypothesis, we measured the elongation rate of frameshifted ribosomes compared to non-frameshifted ribosomes. Fits to the linear portion of run-off curves yielded similar post-tag run-off times of ~386 s and ~475 s, corresponding to average elongation rates of 2.7 ± 0.3 aa/sec and 2.2 ± 0.6 aa/sec, respectively (Fig. 4A and Fig. S7). Fluorescence recovery after photobleaching experiments further confirmed these measurements without the use of drugs (Fig. S8 and Movie S8). Based on these similar elongation rates for both non-frameshifting and frameshifting sites, a single round of translation would take ~9 minutes. Accounting for the number of ribosomes per translation site and their relative fractions, we calculate a cell with 100 RNA would produce ~115 frameshifted protein per hour compared to ~2,220 canonical proteins. In other words, frameshifted proteins would account for ~5% of the total, in agreement with earlier measurements. Thus, the dynamics of the FSS sequence alone can be sufficient to account for the steady-state levels of frameshifted protein in living cells, without the need for additional regulatory mechanisms.

**Figure 4.**
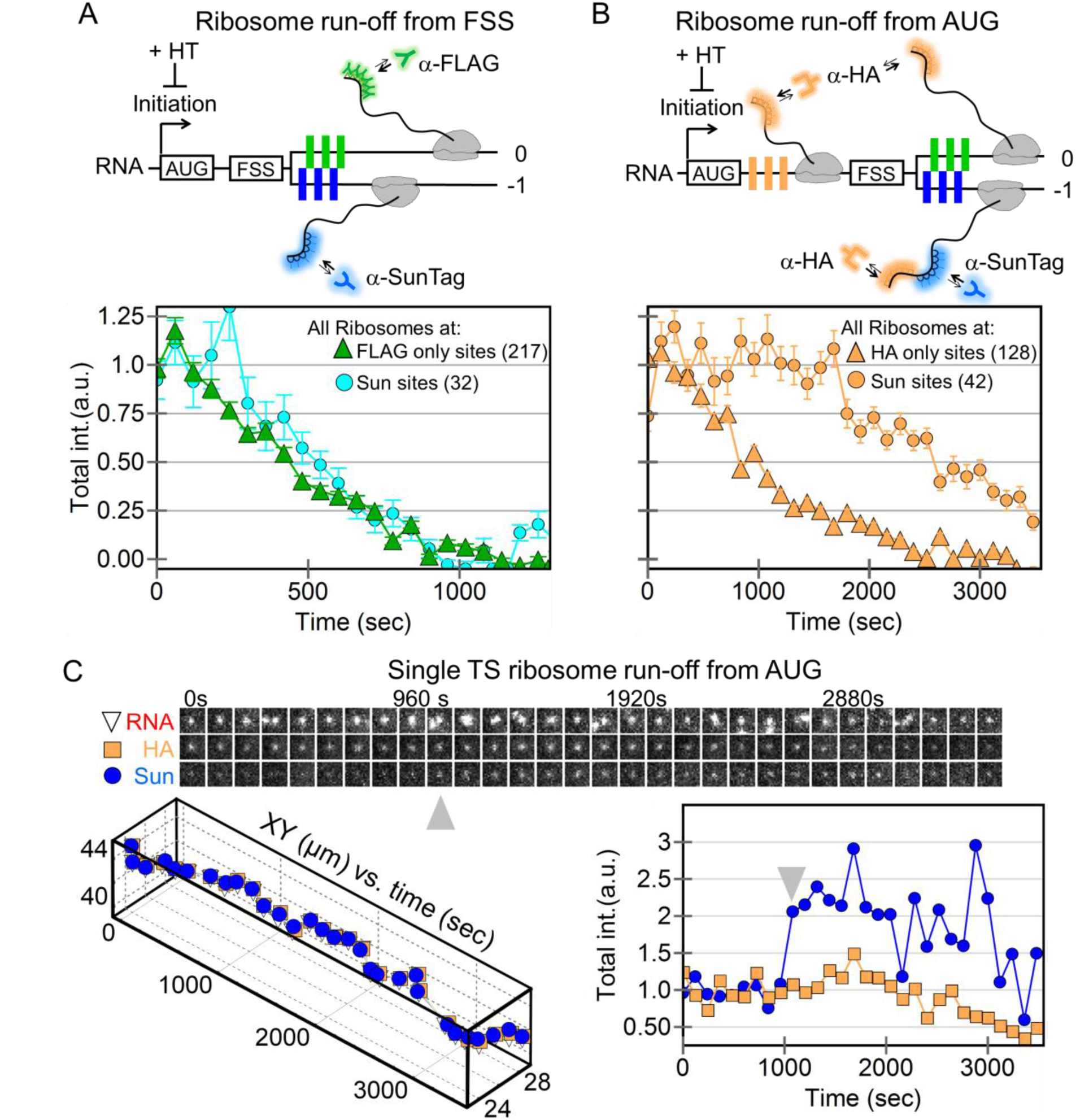
Ribosomal run-off at frameshifting and non-frameshifting translation sites. (A) A schematic showing harringtonine-induced ribosomal run-off from the +FSS 2x multi-frame RNA reporter with FLAG (green) and SunTag epitopes (blue) in the 0 and −1 frames. The intensity of nascent chain signals within non-frameshifting translation sites (marked by just anti-FLAG Fab, green triangles, 217 translation sites initially in 19 cells) and frameshifting sites (marked by anti-SunTag scFv, cyan circles, 32 translation sites initially) drops with time as ribosomes complete translation, i.e. run-off. (B) Same as A, but with a modified multi-frame reporter with extra HA epitopes (orange, HA multi-frame tag) upstream of the frameshift sequence. The intensity of nascent chain signals from non-frameshifting translation sites (marked by just anti-HA Fab, orange triangles, 175 translation sites initially in 27 cells) and frameshifting translation sites (marked by anti-SunTag scFv, orange circles, 42 translation sites initially) drops with time as ribosomes run-off. (C) A sample single translation site encoding the modified HA multi-frame reporter (shown in B) after addition of harringtonine. A montage of image trims shows the detected RNA-, HA-, and Sun-signals through time. Below, the positions of the detected signals within the site are plotted through time. On the right, the normalized total intensity of the HA Fab signal (marking all ribosomes) and the SunTag scFv signal (marking frameshifting ribosomes) is plotted through time. Gray arrows signify a burst of frameshifting, coinciding with multi-RNA interactions.

### Evidence for ribosomal traffic jams at HIV-1 frameshifting translation sites

Despite the similar elongation rates, we noticed a slight (~60 s) delay in the run-off response at frameshifting translation sites (Sun sites Fig 4A). Given earlier work showing the potential of the FSS to pause ribosomes (Dulude et al., 2002), we envisioned this delay could be due to a queue or traffic jam of ribosomes upstream of the FSS within frameshifting sites. As the backed-up ribosomes clear the traffic jam, they replenish the loss of ribosomes running-off. Only after the traffic jam is fully cleared does the number of ribosomes beyond the FSS (with labeled nascent chains) begin to decay.

To test this possibility, we added a 10x HA epitope repeat upstream of the FSS (Fig. 4B). This served two purposes: first, it allowed us to monitor both ribosomes upstream of the FSS (translating HA epitopes) and ribosomes downstream (translating either 0 frame FLAG or −1 frame SunTag epitopes); second, the arrangement more closely mimicked the natural placement of the FSS between the gag-pol polyproteins. In particular, the additional sequence space upstream of the FSS could accommodate longer ribosomal traffic jams, should they occur. We hypothesized that longer ribosomal traffic jams would lead to longer run-off delays. Consistent with this, ribosomes within frameshifting sites took much longer to run-off (Fig. 4B), with frameshifted ribosome levels fluctuating but remaining overall steady for upwards of 3000 s, despite an overall ribosome loss (Figs. S9 and S10). We observed this trend even at the single-molecule level, where the total number of ribosomes (marked by HA) dropped, but the number of frameshifted ribosomes (marked by the SunTag) fluctuated wildly up and down (Fig. 4C, and Movie S9). These fluctuations appear to arise from the stochastic release of stalled ribosomes within the traffic jam and their subsequent translation of frameshifted epitopes. Such a release can be seen in the single-molecule track at the ~1000 s time point, when the frameshift signal gets significantly brighter. Interestingly, this increase in brightness coincided with the association of an additional non-translating +FSS RNA (gray arrows in Fig. 4C). This is consistent with the hypothesis that multi-RNA factories stimulate frameshifting. Although difficult to capture, we observed this type of stimulated frameshifting burst in another single frameshifting RNA track as well (Fig. S11).

### Computational modeling of HIV-1 frameshifting bursts at the single RNA level

To quantify the kinetics of frameshifting, we developed two candidate models and attempted to fit each to our three main observations: (i) the percentages of frameshifting versus non-frameshifting translation sites (Fig. 4C), (ii) ribosomal occupancies (Fig 4D), and (iii) run-off kinetics (Figs. 4E,F). Both models include initiation of ribosomes, codon-dependent elongation of proteins along the RNA template, and ribosomal exclusion to block ribosomes from passing one another or occupying the same place on the RNA. The only difference in the two models is the treatment for how ribosomes shift from the 0 to the −1 frame.

The first model assumes constitutive frameshifting, in which all ribosomes frameshift at the FSS with equal probability. This model could capture either observation (i) or (ii), but not both simultaneously; frameshifting either led to excessively large fractions of frameshifting sites or excessively small ribosomal loading, in disagreement with our observations that a relatively small fraction of RNA frameshift with relatively high ribosomal occupancies. Even with addition of distinct pauses in elongation at the FSS in both frames, the constitutive model was unable to fit our data (Fig. S12).

The second model is inspired by two-state gene models that are commonly used to describe heterogeneous transcription (Munsky et al., 2012). In this `bursty’ model, RNA stochastically switch between non-frameshifting and frameshifting states in which either 0% or 100% of ribosomes produce frameshifted proteins (Fig. 5A). In the bursty model, the RNA frameshift state is assumed to switch ON and OFF at rates *k*_*on*_ and *k*_*off*_, respectively, and the steady state fraction of RNA in the ON state is given by *f* = *k*_*on*_/(*k*_*on*_ + *k*_*off*_). We estimated *f* from the observed fraction of frameshifting translation sites (Fig. 2C).

**Figure 5.**
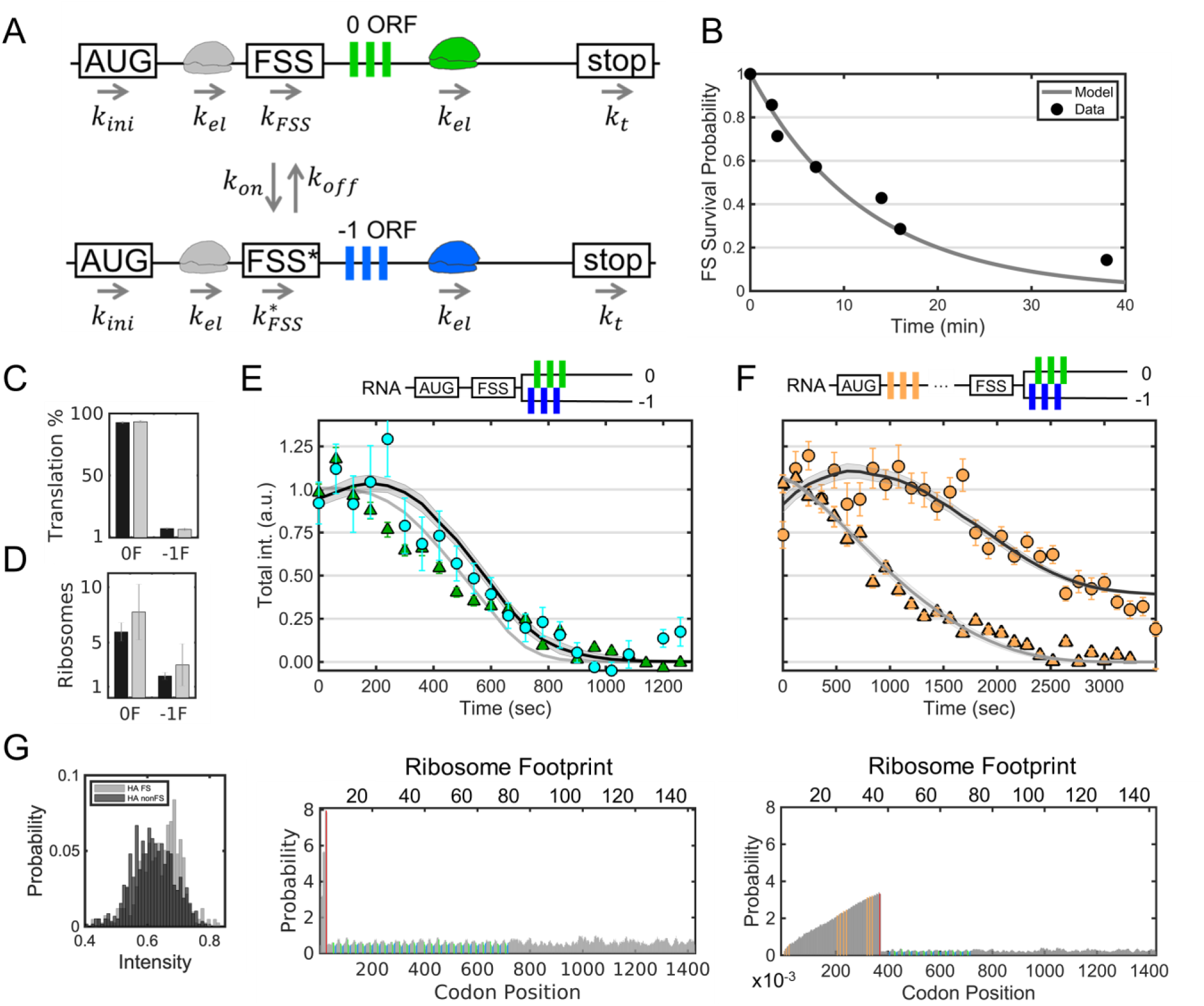
A model for bursty frameshifting. (A) A schematic of the model: *k*_*ini*_ is the translation initiation rate, *k*_*el*_ is the translation elongation rate, *k*_*on*_ is the rate at which RNA switch to the frameshifting state, *k*_*off*_ is the rate at which RNA switch to the non-frameshifting state,*k*_*FSS*_ is the pause rate at the FSS in the non-frameshifting state, 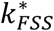 is the pause rate at the FSS in the frameshifting state, and *k*_*t*_ is the termination rate (assumed equal to *k*_*el*_). (B) The survival probability of frameshifting sites through time (gray dots) is fit with a single exponential decay (black line). (C-F) Simultaneous fit of all data. (C) A bar graph comparing the measured (black) and best-fit model predicted (gray) percentage of non-frameshifting (0F) and frameshifting (−1F) translation sites. Error bars represent S.E.M. among cells. (D) A bar graph comparing the measured (black) and best-fit model predicted (gray) number of ribosomes within non-frameshifting (0F) and frameshifting (−1F) translation sites. Error bars represent S.E.M. among RNA. (E) Best-fit model (solid lines) of the data from Figure 4A. Error bars represent S.E.M. among RNA. Below, the predicted ribosomal occupancy along the original multi-frame reporter is shown. The positions of the FSS (red) and FLAG (green) and SunTag (blue) epitopes are shown in color. (F) Best-fit model prediction of the data from Figure 4B. Error bars represent S.E.M. among RNA. Below, the predicted ribosomal occupancy along the modified HA-tagged multi-frame reporter is shown. The positions of the FSS (red) and HA (orange), FLAG (green), and SunTag (blue) epitopes are shown in color. (G) Best-fit model prediction of the intensity distribution of HA signals for frameshifting (gray) and non-frameshifting sites (black) encoding the HA multi-frame tag.

To estimate the timescale of *k*_*on*_ and *k*_*off*_, we tracked translation sites for longer periods of time. To achieve this tracking, we doubled the number of epitopes in the multi-frame reporter (creating the +FSS 2x multi-frame tag) and changed our imaging strategy to sample the RNA signal at all time points and the 0 and −1 translation signals once every fifth time point. This arrangement substantially reduced photobleaching and allowed us to continuously track and monitor the translational status of single translation sites in 3D for nearly an hour. Fig. 5B shows the frameshifting state survival times for the seven translation sites we tracked in this manner, including one site that frameshifted for longer than 40 minutes (Fig. S13 and Movie S10), representing at least four rounds of translation at our measured elongation rate of ~2.4 aa/sec. Remarkably, this frameshifting translation site associated with another for a large part of the 40-minute imaging window, which supports our hypothesis that the FSS facilitates frameshifting in multi-RNA sites. From the frameshift state survival times, we fit the rate of *k*_*off*_ to be ~0.0013 sec^−1^ (Fig. 5B), corresponding to an average frameshift persistence time of 1/*k*_*off*_ ~ 770 s. With *k*_*off*_ and *f* determined, we calculate 1/*k*_*on*_ ~ 10,400 sec. Thus, RNA encoding the HIV-1 frameshift sequence switch to a frameshifting state rarely, on the timescale of a few hours (Table 1, 1/*k*_*on*_). Once an RNA is in the frameshifting state, it remains there for tens of minutes on average (Table 1, 1/*k*_*off*_), occasionally lasting up to an hour or more.

**Table 1:** Bursty Frameshifting Model Parameters.

| **Table 1: Bursty Frameshifting Model Parameters** |  |  |
| --- | --- | --- |
| Ribosome elongation rate $k_{el}$ | 2.4 aa/sec | Simultaneous fit |
| Ribosome initiation rate $k_{ini}$ | 0.02 sec<sup>-1</sup> | Simultaneous fit |
| Switching on rate to frameshifting state $k_{on}$ | 9.6x10<sup>-5</sup> sec<sup>-1</sup> | Simultaneous fit |
| Switching off rate from frameshifting state $k_{off}$ | 1.3x10<sup>-3</sup> sec<sup>-1</sup> | Simultaneous fit |
| Pause rate at FSS in non-frameshifting state $k_{FSS}$ | ~0.0205 aa/sec | Simultaneous fit |
| Pause rate at FSS* in frameshifting state $k_{FSS}^*$ | ~0.011 aa/sec | Simultaneous fit |
| **Calculated quantities with the approximated bursting model assuming HA multi-frame tag** |  |  |
| Fraction of frameshifted RNA | ~6.9% | Calculated |
| Average burst time in the non-shifting state $\langle b \rangle$ | 10,417 sec | Calculated |
| Average burst time in the shifting state $\langle b^* \rangle$ | 770 sec | Calculated |
| Fraction of RNA in in shifting state $f_s$ | 0.069 | Calculated |
| Number of ribosomes to initiate in a shifting state $r_i^*$ | 15 | Calculated |
| Number of ribosomes to initiate in a non-shifting state $r_i$ | 208 | Calculated |
| Time for a ribosome in shifting state to clear the FSS $\tau_{FSS}^*$ | 95 sec | Calculated |
| Time for a ribosome in non-shifting state to clear the FSS $\tau_{FSS}$ | 49 sec | Calculated |
| No. of ribosomes to pass the FSS in the shifted state $r_i^*$ | 8 | Calculated |
| No. of ribosomes to pass the FSS in the non-shifted state $r_c$ | 214 | Calculated |
| No. of ribosomes to accumulate upstream of the FSS in a shifted state $r_a^*$ | 7 | Calculated |
| No. of ribosomes to accumulate upstream of the FSS in a non-shifted state $r_a$ | -5 | Calculated |
| Average elongation time for a ribosome in non-shifting state $\langle \tau_e^* \rangle$ | 645 sec | Calculated |
| Average elongation time for a ribosome in shifting state $\langle \tau_e \rangle$ | 690 sec | Calculated |
| Note: Formulas given in the methods. |  |  |

With these constrained values for *k*_*on*_ and *k*_*off*_, the bursty frameshifting model could reproduce all of our observations when fit by a single set of parameters (Fig. 5C-F and Table 1), in contrast to the constitutive model (Fig. S12). To account for the different run-off delays seen at frameshifting and non-frameshifting sites (seen in Fig. 4A,B), the model required elongation pauses of 1/*k*_*FSS*_ ~ 48 s at non-frameshifting sites and 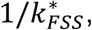 ~ 90 s at frameshifting sites. We explored if codon usage could also explain the differences in run-off times. However, according to the codon adaptation index (Gorgoni et al., 2016; Sharp and Li, 1987), which is related to the speed at which each codon is translated in the simulation, there is no notable difference between the 0 and −1 frames (Fig. S14). Moreover, the distinct pauses predicted by the model are comparable to those previously measured using *in vitro* and *in vivo* bulk assays (Lopinski et al., 2000).

Because the estimated average pause time, 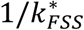, is greater than the average initiation time, 1/*k*_*ini*_, ribosomes could initiate faster than they clear the FSS with an excess rate of 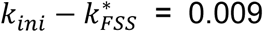 per second and create upstream traffic jams in frameshifting sites. These traffic jams would continue to build for as long as the RNA remains in the frameshifted state, or approximately 1/*k*_*off*_ = 770 s on average. This would allow for the accumulation of approximately 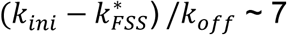 ribosomes (on the HA multi-frame tag, Table 1). Occasionally, traffic jams can extend all the way back to the start codon, as seen in a sample simulation (Movie S11). However, because these traffic jams occur upstream of the FSS within the HA epitope repeats, the nascent chain intensity at translation sites is only partially correlated with ribosomal occupancy. As a result, the model predicts frameshifting sites are slightly brighter than non-frameshifting sites (Fig. 5G), which we also observe experimentally (Fig. S15). Moreover, due to the long time it can take to clear traffic jams, frameshifting can persist for hours following the global shut down of translation initiation (with harringtonine, for example), as can be seen in simulations of the best-fit model (Movie S12) and consistent with what we observed in Figs. 4C and S11. Thus, the final bursty model suggests a mechanism by which frameshifting can persist for long periods in the absence of translation initiation.

## DISCUSSION

Frameshifting is a common tactic used by viruses to minimize their genomes for faster, more efficient replication in host cells, effectively getting two viral proteins for the price of one viral RNA. While the general architecture of frameshift sequences is well characterized and the dynamics of frameshifting have been measured with single-nucleotide precision *in vitro*, until now frameshifting had not been directly observed in a living system. To achieve this, we created a multi-frame repeat epitope tag that can light up single RNA translation sites in different colors depending on which open reading frame is being translated. Together with sensitive single-molecule microscopy and computational modeling, we have demonstrated five novel aspects of HIV-1 frameshifting: (i) frameshifted proteins originate from a small subset of RNA that frameshift with high efficiency; (ii) frameshifting occurs preferentially in multi-RNA translation factories that are facilitated in part by the frameshift sequence; (iii) frameshifting occurs in bursts on single RNA that can last for several rounds of translation; (iv) ribosomes that frameshift are paused for longer at the frameshifting sequence than ribosomes that do not frameshift; and (v) pauses at the frameshift sequence induce ribosomal traffic jams that can maintain the production of frameshifted protein despite global inhibition of translation. Fig. 6 summarizes our findings.

**Figure 6.**
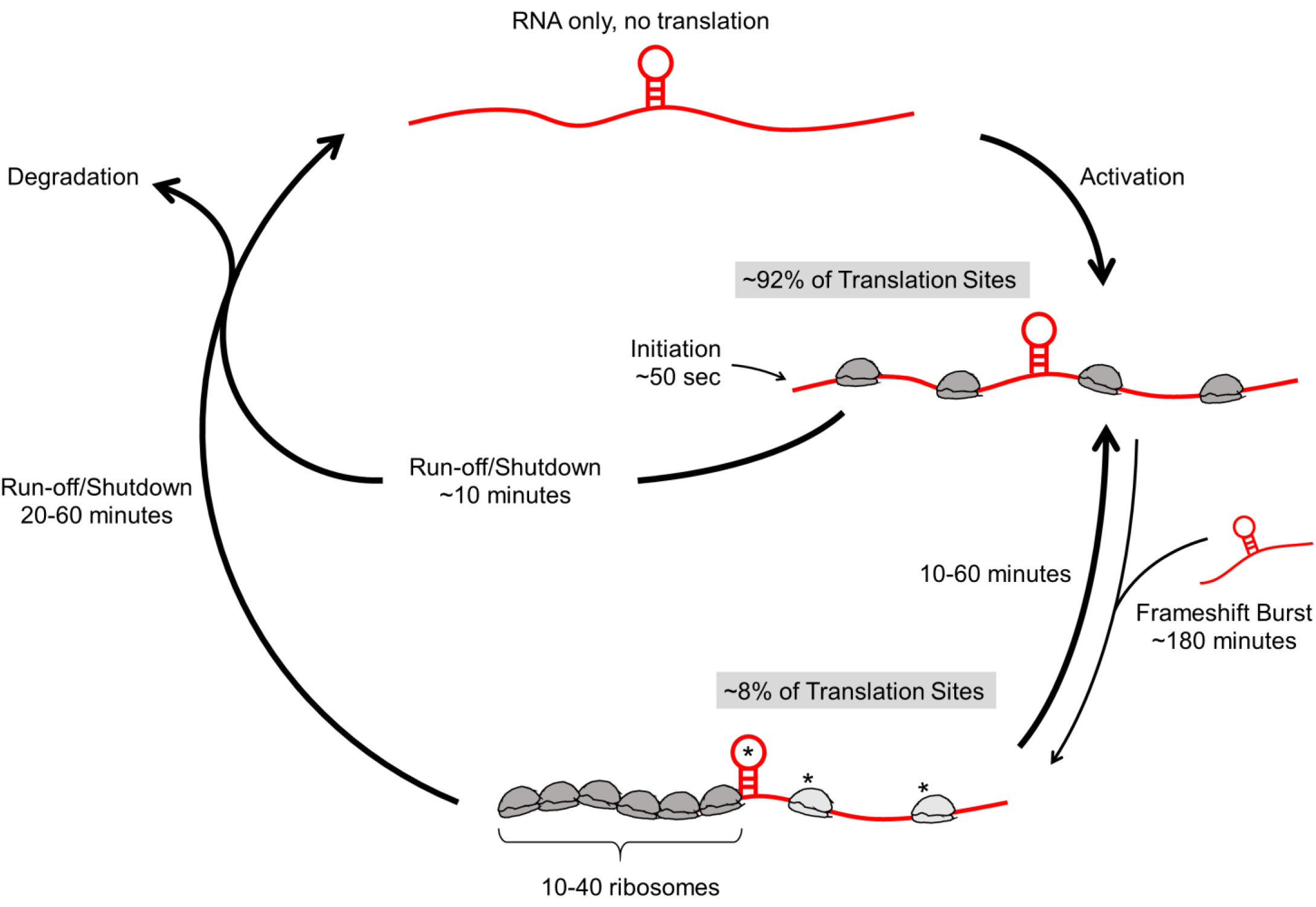
The lifecycle of RNA encoding the HIV-1 frameshift sequence. RNA (red) encoding the HIV-1 frameshift sequence (stem loop) are activated for canonical 0-frame translation. Occasionally, they switch to a frameshifting state (~8%) that can last from tens of minutes to an hour or so. Transitions to this state are stimulated through interactions with other FSS-encoding RNA, either by chance or within multi-RNA translation factories. When in the frameshifting state, longer pauses at the FSS induce longer ribosomal traffic jams. These allow frameshifting RNA to continue to be translated for up to an hour longer than non-frameshifting RNA upon global translation initiation shut-down.

In contrast to constitutive frameshifting on any RNA, our data indicate that frameshifting occurs in bursts on a subset of RNA. Bursty expression has been demonstrated by others, both at single transcription sites as well as at translation sites in bacteria (Lionnet and Singer, 2012) and eukaryotes (Wu et al., 2016). The origin of frameshifting bursts is difficult to pinpoint. It is tempting to speculate that there is a unique structure the RNA takes that enhances frameshifting, particularly given the distinct pause times in our best-fit model. According to our observations, multi-RNA translation factories appear to be more susceptible to frameshift bursts. Such factories have been previously observed (Morisaki et al., 2016; Pichon et al., 2016). Local concentration of translation factors is higher within factories, so translation kinetics are enhanced (Pichon et al., 2016). However, the crowding of machinery at these factory sites may cause them to be more prone to errors and even more so with increased interactions with other FSS-encoding RNA, the FSS being a proverbial wrench within translational factory machinery. Such low-fidelity factories may benefit viral robustness, where mutations are desirable. Consistent with this idea, HIV-1 RNAs prefer a dimeric state which may promote translation factory formation (Barajas et al., 2018).

In this study, we observed frameshifting in the context of factories in three different circumstances. First, in puromycin experiments, we observed frameshifting sites separate into two frameshifting sites upon translation shut-down (Movie S7). Second, we observed on two occasions non-translating RNA interact with a frameshifting site, after which the frameshifting signal at the site markedly increased (Figs. 4E and S11). Third, when we co-transfected FSS oligo RNA into cells (Fig. 3), frameshifting increased in proportion to the amount we co-transfected. Collectively, these data suggest a causal ordering, whereby interactions between FSS-encoding sequences lead to enhanced frameshifting.

According to our best-fit model, pauses always occur at the frameshift sequence, with longer pauses associated with frameshifting RNA compared to non-frameshifting RNA. Pausing in and of itself is therefore only a weak predictor of frameshifting, as others have shown (Kontos et al., 2001; Tu et al., 1992). Nevertheless, the longer pauses associated with frameshifting suggest the frameshift sequence can adopt more than one state or structure, similar to what has been shown *in vitro* with sequences encoding pseudoknots (Houck-Loomis et al., 2011). Longer pauses result in longer ribosomal traffic jams. Our model predicts these jams can extend all the way back to the initiation site, involving up to ~40 ribosomes, as shown in the lower panel of Fig. 5F. Our model predicts an elevated ribosomal occupancy as far back as ~400 codons from the FSS, a length that coincides almost perfectly with the length of the FSS-upstream sequence in HIV-1 gag-pol. This is not unprecedented, as ribosome profiling experiments have also found evidence for relatively high ribosomal densities as far back as the start codon of the gag protein (Napthine et al., 2017). These data suggest the strength of the HIV-1 pause may have evolved to on occasion produce the longest possible traffic jams. As clearance of these jams takes time, frameshifting can persist on RNA for hours despite a global shut-down of translation initiation. In effect, the jam acts like a battery that continually fuels the production of downstream frameshifted protein. This unique mechanism would allow viral proteins to continue to increase in numbers during cellular stress. An open question is how these long traffic jams manage to evade protein quality control (Joazeiro, 2017; Juszkiewicz and Hegde, 2017), which recently was shown to target the interface of jammed ribosomes (Juszkiewicz et al., 2018).

Beyond the imaging of HIV-1 translational frameshifting, the multi-frame tag can now be used in a variety of other contexts. For example, it can immediately be used to investigate frameshifting dynamics along other viral RNA sequences, as well as frameshifting thought to occur along endogenous human mRNA (Cardno et al., 2015). Likewise, the multi-frame tag can be used to examine other non-canonical translation processes involving more than one open reading frame, including start-codon selection, leaky scanning, ribosomal shunting, and general translation fidelity. In fact, in an accompanying paper, a similar multi-frame tag was used to examine upstream and downstream open reading frame selection (Boersma et al.). Like us, the authors also saw a high degree of heterogeneity between translating RNAs, with bursts of translation initiation in multiple open reading frames similar to the bursts of frameshifting we observed at the HIV-1 frameshift sequence. Translational heterogeneity may therefore be far more common than originally appreciated, particularly when it comes to non-canonical translation. We therefore believe multi-color single molecule imaging of both canonical and non-canonical translation will become a powerful new tool for dissecting complex RNA regulatory dynamics in a variety of important contexts.

## Supporting information

## ACKNOWLEDGEMENTS

We thank Dr. Hataichanok (Mam) Scherman of the Colorado State University Protein Expression/Purification Facility and Amanda Koch for cloning and purification of the SunTag scFv-sfGFP and MCP-Halo. We thank all members of the Stasevich and Munsky labs for helpful discussion and comments. This work is supported by the W.M. Keck Foundation.

## AUTHOR CONTRIBUTIONS

Conceptualization, TJS, KL, and BM; Methodology: KL, LA, TM, BM, and TJS; Software, KL, LA, TM, BM, and TJS; Formal Analysis, KL and LA; Experimentation, KL, TM, and TJS; Computational modeling, LA and BM; Resources, BM and TJS; Writing, Review, and Editing, KL, LA, TM, BM, and TJS; Supervision, BM and TJS; Funding Acquisition, BM and TJS.

**SUPPLEMENTARY MOVIE LEGENDS**

**Movie S1. Dynamics of cell presented in Figure 1B**

Max projection of a 13 z-stack movie showing an example U-2 OS cell expressing the +FSS multi-frame tag and bead-loaded with probes: MCP-HaloJF646 (Ch1 red, RNA), FLAG Fab Cy3 (Ch2 green, 0 ORF), and scFv-sfGFP (Ch3 blue, −1 ORF). Images were acquired every 4 seconds for a total of 76 seconds. Circled regions of interest correspond to species identified in Figure 1B and C. Scale bar, 10 µm.

**Movie S2. Dynamics of cells presented in Figure S1**

Single plane movies showing example U-2 OS cells expressing either +FSS multi-frame tag (left), -FSS control multi-frame tag (middle), or -FSS (+1 nt) tag with XXLb1 shifted into the 0 frame (right). All cells were bead-loaded with probes: MCP-HaloJF646 (Ch1 red, RNA), FLAG Fab Cy3 (Ch2 green, AlexX), and scFv-sfGFP (Ch3 blue, XXLb1). Images were acquired every 7 seconds for a total of 42 seconds. Scale bar, 10 µm.

**Movie S3. Dynamics of cell treated with puromycin in Figure 1D**

Max projection of a 13 z-stack movie showing an example U-2 OS cell expressing the +FSS multi-frame tag and bead-loaded with probes: MCP-HaloJF646 (Ch1 red, RNA), FLAG Fab Cy3 (Ch2 green, 0 ORF), and scFv-sfGFP (Ch3 blue, −1 ORF). Images were acquired every 7 seconds for a total of 112 seconds prior to puromycin and 280 seconds following puromycin. Scale bar, 10 µm.

**Movie S4. Dynamics of cell presented in Figure S2**

Max projection of a 13 z-stack movie showing an example U-2 OS cell expressing the +FSS multi-frame tag and bead-loaded with probes: MCP-HaloJF646 (Ch1 red, RNA), FLAG Fab Cy3 (Ch2 green, 0 ORF), and scFv-sfGFP (Ch3 blue, −1 ORF). Images were acquired every 7 seconds for a total of 63 seconds. Circled regions of interest correspond to species identified in Fig. S2. Scale bar, 10 µm.

**Movie S5. Representative cell transfected with the +FSS reverse multi-frame tag**

Max projection of a 13 z-stack movie showing an example U-2 OS cell expressing the +FSS reverse multi-frame tag and bead-loaded with probes: MCP-HaloJF646 (Ch1 red, RNA), FLAG Fab Cy3 (Ch2 green, −1 ORF), and scFv-sfGFP (Ch3 blue, 0 ORF). Images were acquired every 7 seconds for a total of 63 seconds. Circled regions of interest correspond to RNA with no translation (numbered ‘1’), 0 and −1 frame translation site (numbered ‘2’), and 0 frame only translation site (numbered ‘3’). Scale bar, 10 µm.

**Movie S6. Representative cell transfected with the - FSS control tag**

Max projection of a 13 z-stack movie showing an example U-2 OS cell expressing the - FSS control tag and bead-loaded with probes: MCP-HaloJF646 (Ch1 red, RNA), FLAG Fab Cy3 (Ch2 green, 0 ORF), and scFv-sfGFP (Ch3 blue, −1 ORF). Images were acquired every 7 seconds for a total of 63 seconds. Circled regions of interest correspond to RNA with no translation (numbered ‘1’), and 0 frame only translation site (numbered ‘2’-‘8’). Scale bar, 10 µm.

**Movie S7. Zoom-in showing frameshifting site presented as a montage in Figure 1D**

Max projection of a 13 z-stack movie showing an example U-2 OS cell expressing the + FSS multi-frame tag and bead-loaded with probes: MCP-HaloJF646 (Ch1 red, RNA), FLAG Fab Cy3 (Ch2 green, 0 ORF), and scFv-sfGFP (Ch3 blue, −1 ORF). Images were acquired every 7 seconds for a total of 217 (of 280) seconds total post puromycin addition. This movie shows the same cell from Movie S3, but now zoomed in to accentuate a frameshifting site splitting (red arrows). Scale bar, 10 µm.

**Movie S8. Dynamics of Fluorescence Recovery After Photobleaching (FRAP) experiment shown in Figure S8**

Max projection of a 13 z-stack movie showing an example U-2 OS cell expressing the + FSS 2x multi-frame tag and bead-loaded with probes: MCP-HaloJF646 (Ch1 red, RNA), FLAG Fab Cy3 (Ch2 green, 0 ORF), and scFv-sfGFP (Ch3 blue, −1 ORF). Images were acquired every 10 seconds for a total of 100 seconds pre-FRAP and 790 seconds post-FRAP. Control spot (far left red arrow) and two frameshifting sites (center arrows) were tracked through time to measure their fluorescence (0 frame translation) after photobleaching. Scale bar, 10 µm.

**Movie S9. Dynamics of tracked frameshifting site presented in Figure 4C**

Max projection of a 13 z-stack movie showing an example U-2 OS cell expressing the HA multi-frame tag and bead-loaded with probes: MCP-HaloJF646 (Ch1 red, RNA), HA Fab Cy3 (Ch2 green, all ORFs), and scFv-sfGFP (Ch3 blue, −1 ORF). Images were acquired post harringtonine every 120 seconds for a total of 3480 seconds. Scale bar, 10 µm.

**Movie S10. Dynamics of tracked frameshifting sites presented in Figure S13**

Max projection of a 13 z-stack movie showing an example U-2 OS cell expressing the HA multi-frame tag and bead-loaded with probes: MCP-HaloJF646 (Ch1 red, RNA), HA Fab Cy3 (Ch2 green, all ORFs), and scFv-sfGFP (Ch3 blue, −1 ORF). Images shown every 35 seconds for 2275 seconds total. Left is the merge, right is Ch3 only. Scale bar, 10 µm.

**Movie S11. Sample simulation of best-fit bursty model with large ribosomal traffic jam**

A best-fit bursty model simulation of a translation site encoding the HA multi-frame tag. On the top, the simulated intensity of the site is tracked through time, with green showing signal from FLAG epitopes downstream of the FSS, blue showing signal from SunTag epitopes downstream of the FSS, and cyan showing signal from HA epitopes upstream of the FSS. On the bottom, individual ribosomes (gray dots) are shown moving along the reporter (gray line), with green, blue, and cyan squares marking the positions of FLAG, SunTag, and HA epitopes, respectively. As the ribosomes move along the repeat epitopes, the translated nascent chains appear as green (FLAG-tagged nascent chains), blue (Sun-tagged nascent chains), and cyan (HA-tagged nascent chains) dots that increase in size as more epitopes are translated. When the RNA is in the non-frameshifting state, the FSS is labeled ‘FSS(OF)’ in green; when the RNA is in the frameshifting state, the FSS is labeled ‘FSS(−1F)’ in blue. The simulated time is shown to the right of the reporter. A large ribosomal traffic jam is created when the RNA switches into the frameshifting state.

**Movie S12. Sample simulation of best-fit bursty model of frameshifted and non-frameshifted translation sites during harringtonine run-off**

A best-fit bursty model simulation of multiple non-frameshifting (top four labeled ‘nonFS Spots’) and frameshifting (bottom four labeled ‘FS Spots’) translation sites encoding the HA multi-frame tag. In the simulation, harringtonine is added after 1000 s and the simulation continues for an additional 2000 s. Individual ribosomes (gray dots) are shown moving along the reporters (gray lines), with green, blue, and cyan squares marking the positions of FLAG, SunTag, and HA epitopes, respectively. As the ribosomes move along the repeat epitopes, the translated nascent chains appear as green (FLAG-tagged nascent chains), blue (Sun-tagged nascent chains), and cyan (HA-tagged nascent chains) dots that increase in size as more epitopes are translated. When the RNA is in the non-frameshifting state, the FSS is labeled ‘FSS(OF)’ in green; when the RNA is in the frameshifting state, the FSS is labeled ‘FSS(−1F)’ in blue. The simulated time is shown to the right of each reporter.

## Star Methods

### Contact for Reagents and Resource Sharing

Requests for further information and resources should be directed to and will be fulfilled by the corresponding authors, Brian Munsky and Tim

Stasevich. Key plasmids will be deposited on Addgene.

## Materials and Methods

### Plasmid Construction

The HIV-1 frameshift sequence (FSS) followed by either the multi-frame (MF) tag or the reverse multi-frame (revMF) tag were synthesized by GeneArt^®^ gene synthesis service (Thermo Fisher Scientific). The gene fragments were flanked by *NotI* and *NheI*, and fused to the upstream of the beta-actin zipcode and 24x MS2 stem loops in the 3’ UTR of plasmid pUB_smFLAG_ActB_MS2 (Plasmid #81083, addgene) to obtain the FSS-MF and FSS-revMF, respectively. To double the MF tag, the MF tag region was digested out from FSS-MF with *XbaI* and *AgeI*, and then ligated into FSS-MF flanked with *NheI* and *AgeI*. The open reading frame encoding human XXLb1/AlexX (Abramowitz et al., 2004; Aydin et al., 2009) was amplified from U-2 OS cells cDNA with the primers: 5’- GTT GTC ATA TGG GCG TGC GCA ACT −3’; 5’- GAT GTA GCT AGC CTA GAA GCA GCA GGC GGT G −3’. The amplified XXLb1/AlexX was flanked with *NsiI* and *NheI*, and then inserted into the C-terminal region of the FSS-MF, FSS-2xMF and FSS-revMF to obtain FSS-MF- AlexX (i.e. the +FSS multi-frame tag), FSS-2xMF-AlexX (i.e. the +FSS 2x multi-frame tag), and FSS-revMF-AlexX (i.e. the +FSS reverse multi-frame tag), respectively. To produce smHA-FSS-MF-AlexX (i.e. the HA multi-frame tag), the spaghetti monster HA (smHA) (Viswanathan et al., 2015) was flanked with *NotI* and *PstI*, and then inserted into the N-terminal region of FSS-MF-AlexX. For the control constructs, FSS was removed using *KpnI* and *XbaI*. To keep the same frame for MF and AlexX, the following sequence was ligated between *KpnI* and *XbaI* to obtain MF-AlexX (i.e. the -FSS control tag): 5’- GGT ACC GGG AAT TTT CTT CAG AGC AGA CCA GAG CCA ACA GCC GCA CCG TTT CTA GA −3’. To shift the −1 frame into 0 frame for MF and AlexX, the following sequence was ligated between *KpnI* and *XbaI* to obtain MF-AlexX (the -FSS(+1nt) tag): 5’- CGG GAA TTT TCT TCA GAG CAG ACC AGA GCC AAC AGC CGC ACC GTT CT − 3’.

scFv-sfGFP was amplified from pHR-scFv-GCN4-sfGFP-GB1-dWPRE (Plasmid #60907, addgene) using primers: 5’- GCG CGC ATA TGA TGG GCC CCG ACA TC −3’; 5’- GCC GGA ATT CGC CGC CTT CGG TTA CCG TGA AGG T −3’. The amplified scFv-sfGFP was flanked with NdeI and EcoRI, and then inserted into a pET21 vector backbone for expression and purification from *E.coli*.

For HA multi-frame tag experiments, the HA epitope encoded in the scFv-sfGFP plasmid (Plasmid #60907) from Addgene was removed by site-directed mutagenesis with QuikChange Lightning (Agilent Technologies) per manufacturer’s instruction using primers: 5’- CCT CCG CCT CCA CCA GCG TAA TCT GAA CTA GCG GTT CTG CCG CTG CTC ACG GTC ACC AGG GTG CCC −3’; 5’- GGG CAC CCT GGT GAC CGT GAG CAG CGG CAG AAC CGC TAG TTC AGA TTA CGC TGG TGG AGG CGG AGG −3’.

### Fab generation and dye-conjugation

Fab generation was done using the Pierce mouse IgG1 preparation kit (Thermo Fisher Scientific) per the manufacturer’s instructions. Briefly, beads conjugated with ficin were incubated in 25 mM cysteine to digest FLAG (Wako, 012-22384 Anti DYKDDDDK mouse IgG_2b_ monoclonal) or HA (Sigma-Aldrich, H3663 HA-7 IgG_1_ mouse monoclonal) antibodies to generate Fab. Fab were separated from the digested Fc region using a NAb Protein A column (Thermo Scientific, product # 1860592). Fab were concentrated to ~1 mg/ml and conjugated to either Cy3 or Alexa Fluor 488 (A488). Cy3 N-hydroxysuccinimide ester (Invitrogen) or A488 tetrafluorophenyl ester (Invitrogen) was suspended in DMSO and stored at −20°C. 100 µg of Fab were mixed with 10 µL of 1M NaHCO_3_, to a final volume of 100 μL. 2.66 μl of Cy3 (or 5 μl of A488) was added to this 100 μL mixture and incubated for 2 hours at room temperature with end-over-end rotation. The dye conjugated Fab were eluted from a PBS equilibrated PD-mini G-25 desalting column (GE Healthcare) to remove unconjugated dye. Dye conjugated Fabs then were concentrated in an Ultrafree 0.5 filter (10k-cut off; Millipore) to 1 mg/ml. This conjugation and concentration process was repeated on occasion to ensure a degree of labeling close to one. The ratio of Fab:dye, *A*_*rat*_, was determined using the absorbance at 280 and 550 nm or 495 nm, the extinction coefficient of IgG at 280 nm, Ɛ_*IgG*_, the extinction coefficient of the dye, Ɛ_*dye*_, provided by the manufacturer, and the dye correction factor at 280 nm, *CF*, provided by the manufacturer. The degree of labeling, *DOL*, was calculated with the following formula:

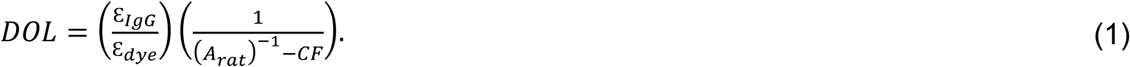

Only Fab calculated with a *DOL* ~1 were used in experiments.

### MCP and scFv-sfGFP purification

His-tagged MCP or scFv-sfGFP was purified over a Ni-NTA-agarose (Qiagen) per the manufacturer’s instructions with minor modifications. Briefly, bacteria were lysed in a PBS-based buffer with a complete set of protease inhibitors (Roche). Binding to the Ni-NTA resin was done in the presence of 10 mM imidazole. The resin was washed with 20 and 50 mM imidazole in PBS. The protein was then eluted in 300 mM imidazole in PBS. The eluted his-tagged MCP was dialyzed in a HEPES-based buffer (10% glycerol, 25 mM HEPES pH 7.9, 12.5 mM MgCl_2_, 100 mM KCl, 0.1 mM EDTA, 0.01 % NP-40 detergent, and 1 mM DTT), snap-frozen in liquid nitrogen, and stored at −80C.

### Cell culture, transfection, and bead-loading

U-2 OS cells were grown using DMEM (Thermo Scientific) supplemented with: 10% (v/v) FBS, 1 mM L-glutamine and 1% (v/v) Penicillin-streptomycin. Before experiments, cells were plated on a 35 mm MatTek chamber (MatTek) and DNA was either transiently transfected with Lipofectamine LTX (Thermo Scientific) per the manufacturer’s instructions or transiently transfected via bead-loading. As described previously (Hayashi-Takanaka et al., 2011; Morisaki et al., 2016), bead-loading involved the following six steps: First, 100 μg/ml of fluorescently labeled Fab, 250 μg/ml of purified (GCN4) scFv-sfGFP and 33 μg/ml of purified MCP-HaloTag protein were mixed with PBS to a final volume of 4 μl. Second, in cell culture hood, DMEM was aspirated from the MatTek chamber and the 4 μl mix was pipetted on top of cells. Third, ~106 μm glass beads (Sigma Aldrich) were evenly sprinkled over the cells. Fourth, the chamber was tapped carefully ~10 times on the cell culture hood bench top. Fifth, DMEM was immediately added back to the cells. Sixth, cells were returned to the incubator for at least an hour to recover from the loading procedure. In most experiments, we also bead-loaded DNA, which we added to the initial 4 μl mix (so that DNA had a final concentration close to 1 mg/ml). On occasion, DNA was transiently transfected ~2 hours before bead-loading. Around one hour before experiments began, bead-loaded cells were washed with phenol-red-free complete DMEM to remove glass beads, and 200 nM of JF646-HaloTag ligand (a cell permeable fluorogenic ligand (Grimm et al., 2015)) was added to label MCP-HaloTag protein. After 30 mins of incubation, cells were washed three times using phenol-red-free complete DMEM to remove any unconjugated fluorophores. Cells were then immediately imaged for experiments.

### Single molecule tracking microscopy

To track single molecule translation sites, a custom-built widefield fluorescence microscope based on a highly inclined illumination scheme (Tokunaga et al., 2008) was used (Morisaki et al., 2016). Briefly, the excitation beams, 488, 561 and 637 nm solid-state lasers (Vortran), were coupled and focused at the back focal plane of the objective lens (60X, NA 1.49 oil immersion objective, Olympus). The emission signals were split by an imaging grade, ultra-flat dichroic mirror (T660lpxr, Chroma) and detected using two EM-CCD (iXon Ultra 888, Andor) cameras via focusing with 300 mm tube lenses (producing 100X images with 130 nm/pixel). With this setting, one camera detected far-red signals and the other detected either red or green signals. Far red signals were detected with the 637 nm laser and the 731/137 nm emission filter (FF01-731/137/25, Semrock). Red and green signals were separated by the combination of the excitation lasers and the emission filters installed in a filter wheel (HS-625 HSFW TTL, Finger Lakes Instrumentation); namely, the 561 nm laser and 593/46 nm emission filter (FF01-593/46- 25, Semrock) were used for Cy3 imaging, and the 488 nm laser and 510/42 nm emission filter (FF01-510/42-25, Semrock) were used for sfGFP or A488 imaging. Live cells were placed into a stage top incubator set to a temperature of 37°C and supplemented with 5% CO_2_ (Okolab) on a piezoelectric stage (PZU-2150, Applied Scientific Instrumentation). The focus was maintained using the CRISP Autofocus System (CRISP-890, Applied Scientific Instrumentation). The lasers, the cameras, the filter wheel, and the piezoelectric stage were synchronized via an Arduino Mega board (Arduino). Image acquisition was done with open source Micro-Manager software (Edelstein et al., 2014). Imaging size was set to 512 × 512 pixels^2^ (66.6 × 66.6 μm^2^), and exposure time was set to 53.64 msec. Readout time for the cameras from the combination of our imaging size, readout mode, and the vertical shift speed was 23.36 msec, resulting in an imaging rate of 13 Hz (70 msec per image). The excitation laser lines were digitally synched to ensure they only illuminated cells when the camera was exposing in order to avoid excessive photobleaching. To capture the entire volume of the cytoplasm of U-2 OS cells, 13 z-stacks with step size of 500 nm (6 μm in total) were imaged using the piezoelectric stage such that the z-position changed every 2 images (one image for Cy3 and one for sfGFP/A488 + JF646). The position of the filter wheel was changed during the camera readout time. This resulted in a total cellular imaging rate of 0.5 Hz (2 sec per volume for 3-colors). Note that all colors described in the text and that are shown in the figures are based on the color of the excitation laser: RNA in red (JF646) and protein in green (Cy3) or blue (sfGFP/A488).

### Particle tracking

Images were first pre-processed using either Fiji (Schindelin et al., 2012) or a custom-written batch processing *Mathematica* code (Wolfram Research) to make 2D maximum intensity projections from 3D images. Pre-processed images were then analyzed with a custom-written *Mathematica* code to detect and track particles. Specifically, particles were emphasized with a band-pass filter so the positions could be detected using the built-in Mathmatica routine ComponentMeasurements “IntensityCentroid”. Detected particles were linked through time by allowing a maximum displacement of 5 pixels between consecutive frames. Particle tracks lasting at least 5-10 frames were selected and their precise coordinates were determined by fitting (using the built-in *Mathematica* routine NonlinearModelFit) the original 2D maximum intenstiy projected images to a 2D Gaussians of the following form:

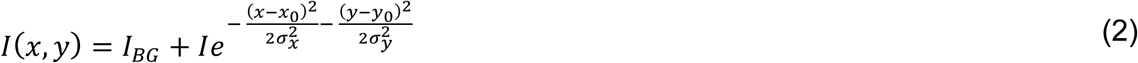

where *I*_*BG*_ is the background intensity, *I* the particle peak intensity, (σ_*x*_, σ_*y*_) the spread of the particle, and (*x*_0_, *y*_0_) the particle location. The offset between the two cameras was registered using the built-in *Mathematica* routine FindGeometricTransform to find the transform function that best aligned the fitted positions of 100 nm diameter Tetraspeck beads evenly spread out across the image field-of-view. We did not register the images, but only the fitted positions in order to avoid introducing any distortion into images. This is why a slight offset can be observed between the red and the green/blue particles even though they are within a diffraction limited spot, according to our registration.

For visuallization and some quantification, average intensity image trims were created by averaging images of all detected particles of a given species (each centered by their intensity centroid). To compute the average number of ribosomes in a translation site (see Fig. S4), the average image trims of translation sites and single mature proteins were fitted to Eq. (2). The average intensities were calculated by integrating the fitted Gaussian. The ratio of the average intensity of translation sites to that of single mature proteins was then calculated to give the average number of ribosomes in a translation site. The average intensity of single RNAs were also calculated using the same procedure (Fig. 3C, S5, and S6). Fits were susceptible to noise, so we also used an alternative strategy to determine the average intensity of translation sites and RNA that was robust to noise (in all Figs aside from Fig. 3C, S4, S5, and S6). Specifically, the intensity of centered images of RNA or translation sites was calculated to be the average intensity within a centered radial spot of four pixels in diameter minus the average background intensity from a centered ring with an outer diameter of twelve pixels and an inner diameter of eight pixels. *Mathematica* source code is available upon request.

### Translation Site Species Identification

After RNA particles were identified and tracked using the custom *Mathematica* code described above, an average centered image of the first five frames from each track was created for RNA (JF646), 0 ORF (FLAG-Cy3 in the +FSS multi-frame tag), and −1 ORF (scFv-sfGFP in the +FSS multi-frame tag). The trims were then hand checked to remove any trims with artifacts, e.g., smears or non-diffraction limited spots. Next, a custom *Mathematica* code was used to detect particles in the 0 ORF or −1 ORF trim channels, sorting the spots into RNA only, 0 frame translation sites (0 TS only), 0 and −1 TS, and − 1 only TS. For all cases, RNA always had to be present. Finally, frameshifting translation sites (the 0 and −1 TS or the −1 only TS) were validated by eye, to further remove artifacts. For example, RNA that briefly colocalized with a mature protein puncta were removed at this stage. After all sites were validated, the total count of each type of species was used to determine the percentage of non-translating RNA (no TS), 0 frame translation sites (0 TS), 0 and −1 TS, and −1 only TS.

### Puromycin treatment

To confirm active translation elongation, puromycin (Sigma Aldrich) was used to release nascent chains from elongating ribosomes, leading to a rapid loss of nascent chain signal at translation sites. Bead-loaded cells with visible translation sites were imaged at a rate of one volume every seven seconds. After acquiring 16 pre-images, cells were treated with a final concentration of 100 μg/ml puromycin and continuously imaged for an additional 100 time points. As a control, the same imaging conditions were performed except that the cells were treated with vehicle (H_2_O). In this case, nascent chain signals did not disappear (data not shown). Both frameshifting and non-frameshifting translation sites were monitored through time using the tracking code described above.

### Oligo FSS RNA co-transfection

An uncapped RNA oligo containing the FSS from HIV-1 was synthesized from IDTDNA with the following RNA sequence: 5’- CUG GCC UUC CCU UGU GGG AAG GCC AGA UCU UCC CUA AAA AA −3’. Co-transfection of FSS-MF-AlexX and the RNA oligo was carried out via bead-loading, as describe above. Briefly, 1 μg plasmid FSS-MF-AlexX construct DNA, 100 μg/ml of fluorescently labeled Fab, 250 μg/ml of purified (GCN4) scFv-sfGFP or 1 μg of (GCN4) scFv-sfGFP plasmid, 1 or 4 μg of RNA oligo, and 33 μg/ml of purified MCP-HaloTag protein were mixed with PBS to a final volume of ~4 μl of PBS.

### Ribosome Run-Off Experiments and fits

To measure the average elongation rate, harringtonine (Cayman Chemical) was used to block translation initiation and iduce the run-off of all actively elongating ribosomes, leading to a gradual loss of nascent chain signal at translation sites. Beadloaded cells with visible translation sites were imaged as described above except that cell volumes were acquired every 60 or 120 seconds. After acquiring 5 pre-images, cells were treated with a final concentration of 3 μg/ml Harringtonine and continuously imaged for 30 more time points. As a control, the same imaging conditions were performed except that cells were treated with vehicle (DMSO). In this case, nascent chain signals did not decay (data not shown). The intensities of translation sites measured, as described above. The intensity of all translation sites in each frame (and in all cells) were then totaled to produce the run-off curve. Run-off curves were normalized to the mean of the total intensity of the first four time points after treatment of harringtonine began. These curves were then fit to a linear regression to estimate run-off times (see Fig. S7). The linear portion of the run-off decay begins when the normalized run-off intensity reaches a fraction *f*_0_, where

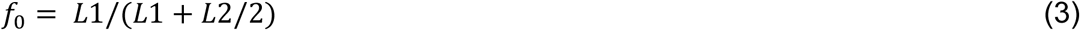

where *L*1 is the length of untagged portion of the open reading frame and *L*2 is the length of the tagged portion (i.e. the length of the repeated epitopes), as described (Morisaki and Stasevich, 2018). The linear portion of the decay was then interpolated to background levels to estimate the run-off time *R*_*T*_. The elongation rate *k*_*el*_ was calculated as follows:

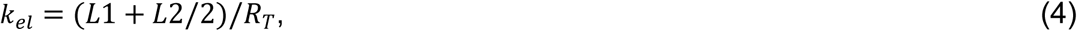

### Fluorescence recovery after photobleaching (FRAP)

To confirm elongation rates of frameshifting translation sites, FRAP experiments were performed. Bead-loaded cells with visible translation sites were imaged once every 10 seconds, with an intentional photobleach at frame 10. The photobleach was performed with a 405 nm laser focused to a spot roughly 5 μm in diameter and operating at a minimal power such that the nascent chain signal did not completely vanish. This allowed us to track the translation site continously throughout the experiment. Following the intentional photobleach, the fluorescence recovery of translation sites within the bleach zone were monitored for an additional 80 time points. These translation sites were tracked and their intensities quantified, as described above. To correct for unintentional photobleaching, the intensity of a control translation site was also tracked and quantified. The loss of signal from the control translation site was fit to a single exponential decay and this decay was divided out from the FRAP recovery curves of the intentionally photobleached translation sites. The FRAP recovery curve can be thought of as the inverse of the harringtonine run-off curve (Morisaki and Stasevich, 2018). In this way, the FRAP recovery curve was fit to determine the average translation elongation rate (Fig. S8).

### General aspects of modeling frameshift kinetics

The frameshift models describe the stochastic dynamics of nascent translation with single-codon resolution. Both the constitutive and bursting models are formulated to track an arbitrary number of individual ribosomes that can perform three possible reactions: (i) A new ribosome can initiate translation at the start codon at rate *k*_*ini*_. (ii) An existing ribosome can elongate at rate *k*_*e*_ to incorporate one amino acid into the nascent peptide chain. The rate *k*_*e*_ is sequence-specific with each codon’s rate scaled by the genomic copy of the corresponding codon (Nakamura et al., 1999). When the ribosome completes the final amino acid, translation is terminated and that ribosome is eliminated. (iii) A ribosome at the frameshift sequence can shift from the 0 to the −1 frame. In addition to these three reactions, ribosomes at the frameshift sequence can pause for an average time of 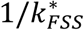 or 1/*k*_*FSS*_ for the shifted and non-shifted states. For the bursting model, each RNA is assumed to switch back and forth between non-frameshifting and frameshifting states with rates *k*_*on*_ and *k*_*off*_. With these mechanisms and parameters, the models can be analyzed using either simplified approximations or detailed simulations.

Several approximate features can be derived directly from the bursting model parameters (see Table 1). The average burst time or time spent in a frameshifting state is:

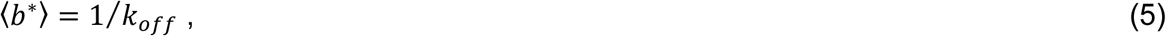

The average time spent in a non-frameshifting state is:

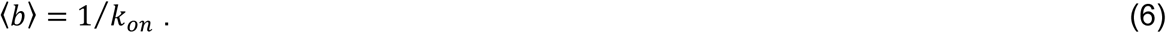

The fraction of RNA in a frameshifting state is:

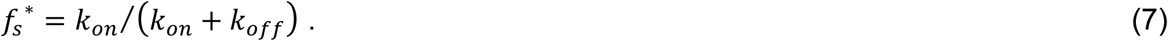

The number of ribosomes that initiate during the frameshifting state is:

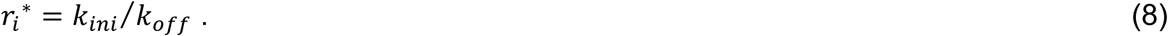

The number of ribosomes that initiate during the non-frameshifting state is:

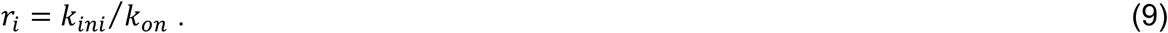

The time for a ribosome to clear the FSS in the frameshifting state is:

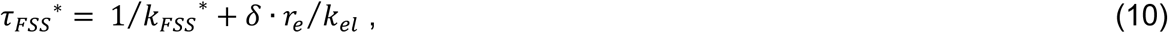

where the second term accounts for ribosome pileup in the 9-codon ribosomal exclusion (Ingolia et al., 2009), *r*_*e*_, region upstream from the FSS. Similarly, the time for a ribosome to clear the FSS in the non-frameshifting state is:

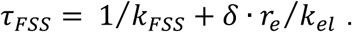

One can approximate *δ* = 1 or 0 if initiation is faster (*k*_*ini*_ > *k*_*FSS*_) or slower (*k*_*FSS*_ > *k*_*ini*_) than the FSS pause. The number of ribosomes to pass the FSS in the frameshifting state is:

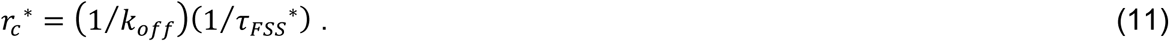

The number of ribosomes to pass the FSS in the non-frameshifting state is:

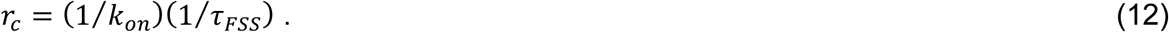

The average number of ribosomes to accumulate upstream of the FSS in the frameshifting state is:

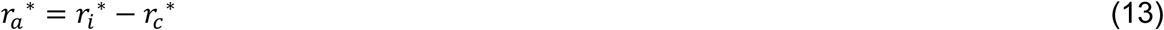

and the number to clear in a non-frameshifting state is:

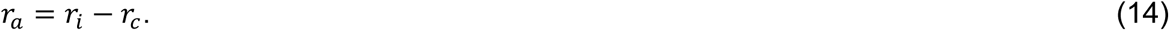

In equations (13) and (14), positive values of *r*_*a*_ correspond to accumulation of ribosomes in the traffic jam, and negative values correspond to clearance of ribosomes. The average elongation time for a ribosome in the frameshifting state is:

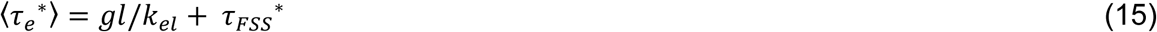

and the number to clear in a non-frameshifting state is:

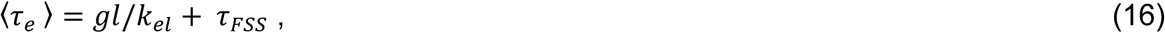

where *gl* is the gene length in codons.

Simulations were started at *t* = −10,000 seconds to approximate steady state at t = 0 using the Gillespie algorithm (Gillespie, 1976). Ribosome densities were found by collecting position statistics for multiple simulations. Simulated ribosome numbers and positions and multi-frame tag probe locations were combined to estimate translation site intensities. Harringtonine assays were simulated by preventing the initiation reaction at the time of treatment. Parameter estimation was performed using genetic algorithms and a multiple-objective cost function that considers the frameshifting efficiency, the number of ribosomes per RNA and the Harringtonine assays. A detailed description of the computational methods and codes is given in the ‘Computational Details’ section below.

### Computational Details

Both the constitutive model and the bursting model consist of three general ribosomal reaction types: initiation (*w*_0_), elongation (*w*_*i*_ or 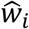) and frameshifting (*w*_*FSS*_ or 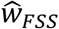) as depicted in Eq. 17,

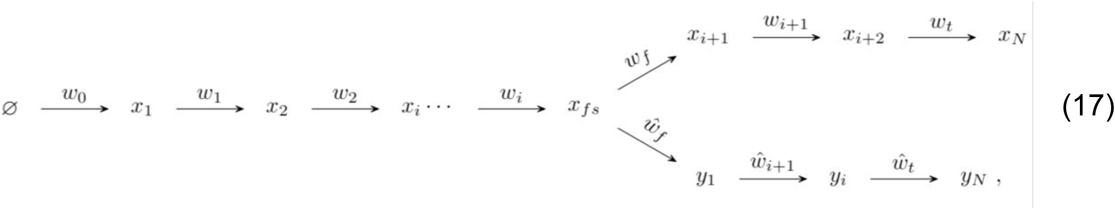

#### Ribosome Initiation

Ribosomes are assumed to bind to the RNA with rate *k*_*ini*_. To account for the fact that ribosomes are large biomolecules that occupy around 20 to 30 nuclear bases on the RNA (Ingolia et. al, 2009), initiation is blocked by any downstream ribosome within *n*_*f*_= 9 codons (in either frame). Therefore, initiation rate is set at:

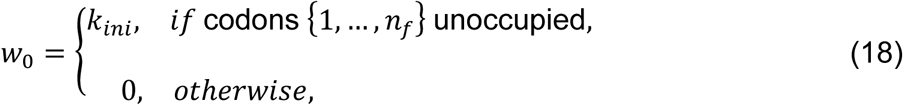

#### Ribosome Elongation

Each ribosome moves along the RNA codon by codon in the 5’ to 3’ direction. The elongation rate for each *i*^*t*ℎ^ codon, 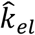 is assumed to be:

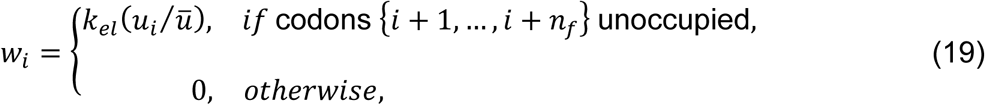

where *u*_*i*_ denotes the codon usage frequency in the human genome obtained from (Nakamura et. al, 2000), and 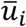 represents the average codon usage frequency in the human genome. The fit parameter *k*_*el*_ specifies the average elongation rate. Ribosomal termination is assumed to be equivalent to elongation of the final codon.

#### Frameshifting and Pausing

When the ribosome reaches the frameshift site, *n*_*FSS*_, it pauses, and may shift from the 0 frame to the −1 frame. For the constitutive model, the ribosome can continue in the 0 frame with rate

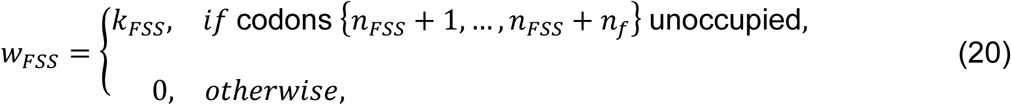

or the ribosome can continue in the −1 frame at rate:

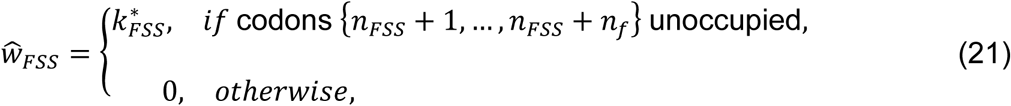

For the bursting model, the decision to continue in the 0 or −1 frame depends upon the frameshifting state with the rate given by:

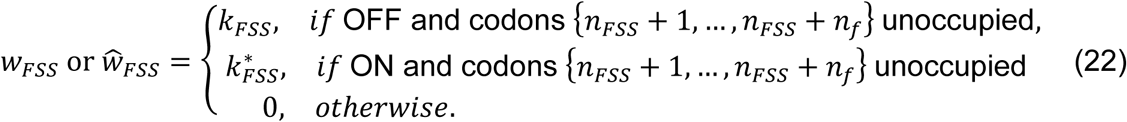

#### Relating model dynamics to experimental fluorescence intensity

To relate ribosome occupancy to experimental translation spots, we proscribe an assumed fluorescence to each ribosome based upon its position. This intensity is proportional to the number of peptide probes upstream from the ribosome location and in the appropriate frame or frames. Ribosomes in the 0 frame include all upstream probes in the 0 frame. Ribosomes in the −1 frame include probes in the 0 frame between the start codon and the FSS (if any) and probes in the −1 frame between the FSS and the current ribosome position. The total intensity vector for the *j*^*th*^ color is given by

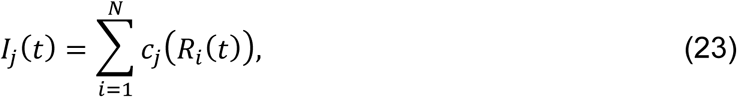

where *R*_*i*_(*t*) denotes the position (i.e., frame and codon location) of the *i*^*th*^ ribosome, and *c*_*j*_(*R*_*i*_) is the corresponding intensity of the *j*^*th*^ color.

### Comparison of Data and Models

#### Fraction of translation spots

Experimental data was measured between 2 to 6 hours post bead-loading for a period of 120 seconds. To reproduce these experimental data, the model was solved using 100 trajectories of 120 seconds starting at steady state. Spots were classified as 0 frame only (*s*_0*F*_), −1 frame only (*s*_−1*F*_) and both frames (*s*_*BF*_). Following the experimental observation that frameshifting spots commonly consist of more than one RNA, *s*_*BF*_ were equally added to the *s*_0*F*_ and *s*_−1*F*_. The percentage of non-frameshifting translation spots was calculated as follows:

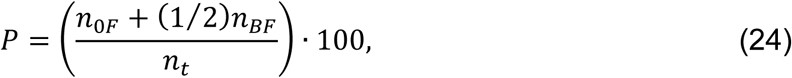

where *n*_0*F*_ is the sum of *s*_0*F*_ and *n*_*t*_ is the total number of translating spots.

The percentage of frameshifting translation spots was calculated as follows:

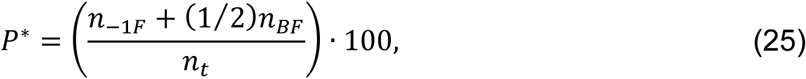

where *n*_−1*F*_ is the sum of *s*_−1*F*_.

#### Number of ribosomes per RNA

Experimental observations allowed to quantify the number of ribosomes in frameshifting and non-frameshifting translation spots (Figure 2 in the main text). To relate model simulations and data, the number of ribosomes per RNA was estimated by directly by adding the number of ribosomes in each RNA at steady state.

#### Harringtonine assays

Harringtonine inhibits translation by binding to the ribosomal 60S sub-unit, which blocks new initiation events. Experimental data showed that Harringtonine causes the intensity in translating spots to drop to a basal intensity value after a run-off time (Figure 4 in main text). To mimic the effects of Harringtonine in our model we modified the initiation rate as:

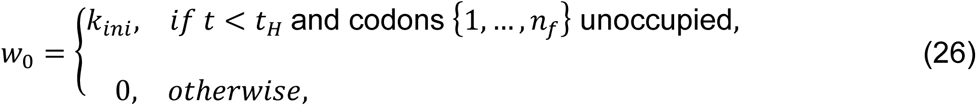

where *t*_*H*_ is the time of application of Harringtonine. After Harringtonine application, spots simulated from the original construct were classified as 0 frame only (‘0F’) or both 0 frame and −1 frames (‘BF’). After classification, average spot intensities were quantified as:

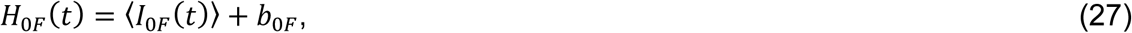

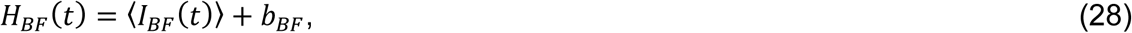

where *b*_(.)_ is experimental background expression obtained at the end of the experimental time of the run-off assays. Similarly, spots simulated for the extended construct with upstream HA tags and downstream −1 tags were classified as HA in non-shifted spots (‘HA’) or HA in shifted spots (‘HA*’).

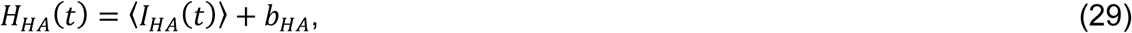

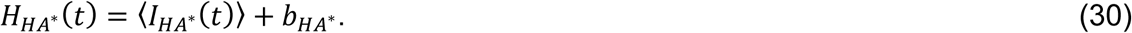

#### Parameter estimation

For each model, we sought to find a single parameter set that reproduces all experimental data. Given the different sources of experimental data we estimated the parameter values using a multi-objective optimization strategy to simultaneously compare the Harringtonine run-off data, the fractions of shifted and non-shifted spots and the ribosome occupancy, as follows:

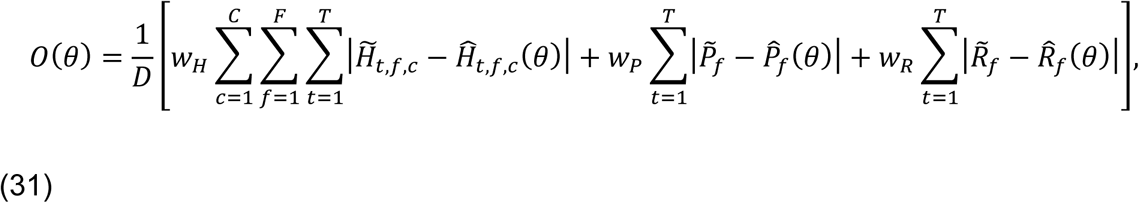

where experimental data are denoted as 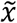 and simulations results as 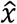. Weights (*w*_*H*_, *w*_*P*_ and *w*_*R*_) were used to balance constraints by the different experiments and time points. *T* represents the number of experimental points in the Harringtonine assays (*T* = 30), *F* is the number of studied frames (*F* = 2), *C* is the number of different gene constructs (*C* = 2), and *D* is the number of different data types (*D* = 3). The weight used on the Harringtonine data set was defined as *w*_*H*_ = 1⁄(*C* ∙ *F* ∙ *T*). The weight in the fraction of spots per frame was defined as *w*_*P*_ = 1⁄(*P* + *P*^∗^). And the weight for the number of ribosomes was defined as: *w*_*R*_ = 1⁄(*R* + *R*^∗^).

#### Parameter searches

Parameter searches consisted of optimization routines based on genetic algorithms (GA) using a population of 100 individuals and running the search for 20 generations.

The constitutive model has a total of four parameters (*k*_*el*_, *k*_*ini*_, *k*_*FSS*_ and 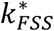), whereas the bursting model has two additional parameters (*k*_*on*_, *k*_*off*_). The optimization strategy consisted in to fit four free parameters (*k*_*el*_, *k*_*ini*_, *k*_*FSS*_ and 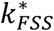) in the models, parameters (*k*_*on*_, *k*_*off*_) were independently determined from data in Fig. 5B,C). The parameter set that best reproduces the data was selected as:

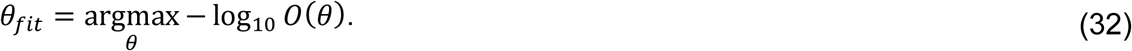

Optimized parameter values are given in full detail in table 1 for the bursting model.

#### Simulation details

To simulated the model’s stochastic dynamics, we used the direct method from Gillespie’s algorithm (Gillespie 1976) coded in Matlab. Genetic Algorithms were coded in Matlab. Simulations were performed on the W. M. Keck High Performance Computing Cluster at Colorado State University.

---

**Fig. S1.**
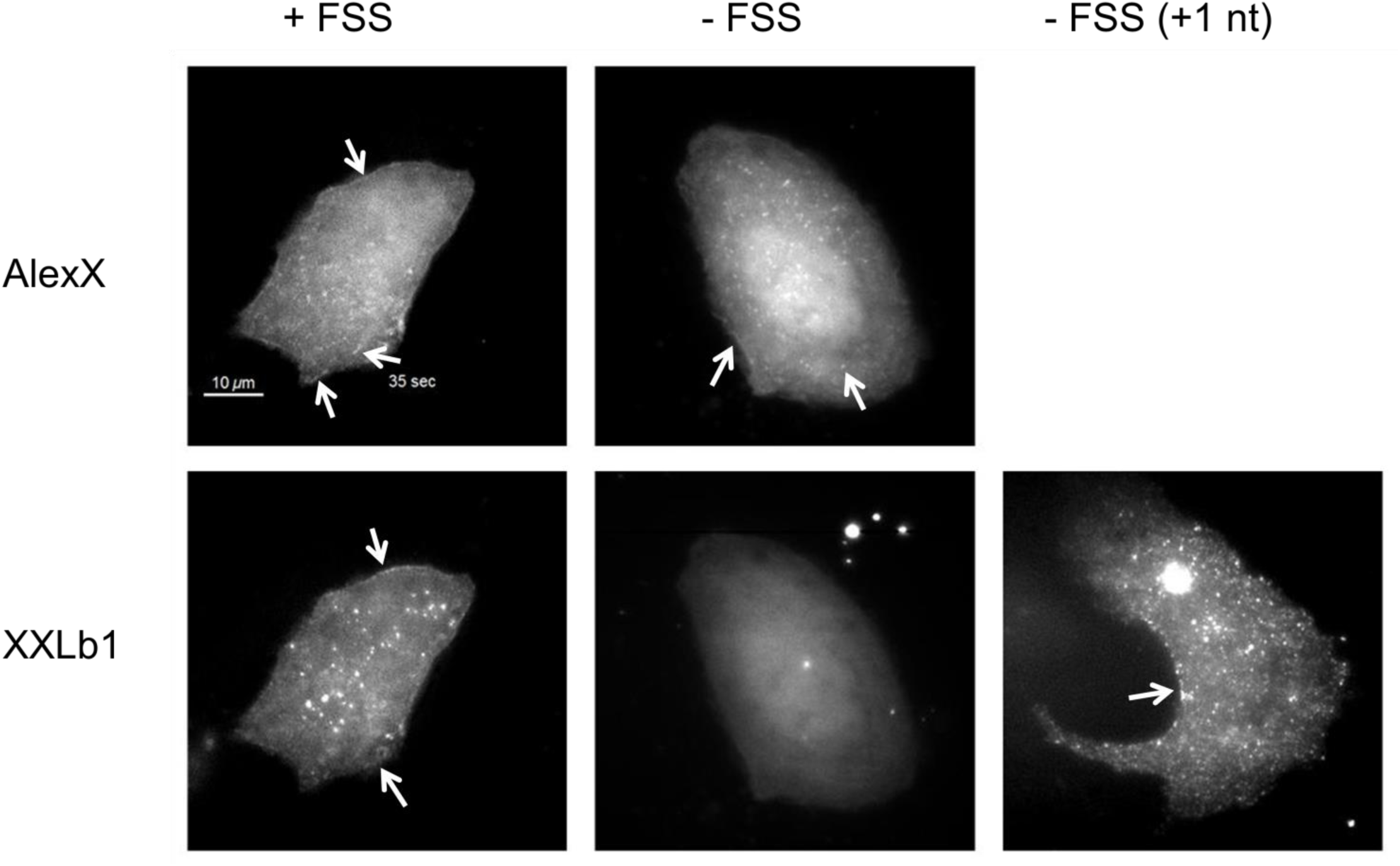
Accumulation of mature AlexX and XXLb1 proteins from multi-frame tags. Mature AlexX (upper) and XXLb1 (lower) membrane proteins accumulate in distinct punctae that coalesce to form elongated structures that undulate dynamically. These structures are enriched at the top and bottom of cells and around the edge of the cytoplasm (arrows), consistent with membrane localization. Movie S2 shows the dynamics of these structures. In the +FSS multi-frame reporter, mature AlexX comes from 0 frame canonical translation, while mature XXLb1 comes from −1 frameshifting translation. In the -FSS control reporter, only mature AlexX accumulates, indicating little to no frameshifting. The -FSS (+1 nt) column shows accumulation of XXLb1 when an extra nucleotide is inserted in the -FSS control reporter following the start codon. This extra nucleotide pushes XXLb1 into the 0 frame, demonstrating it can be expressed.

**Fig. S2.**
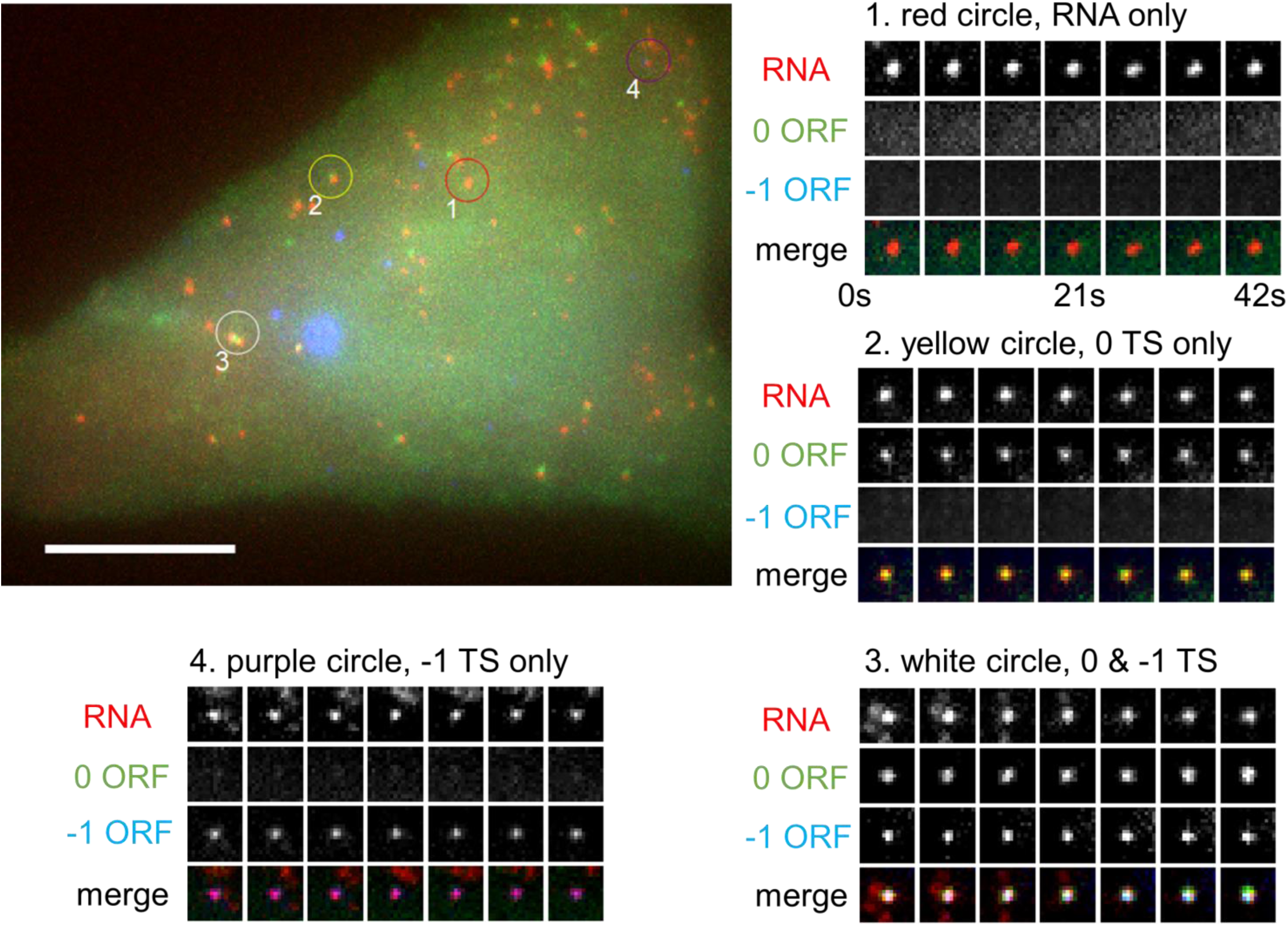
A cell with all possible types of translation sites. A sample cell exhibiting all four possible translation sites (TS). (1) untranslated TS; (2) 0 frame only TS; (3) 0 and −1 frame translated TS; and (4) −1 frame only TS. Scale bar = 10 μm. Movie S4 showcases the dynamics of these spots.

**Figure S3.**
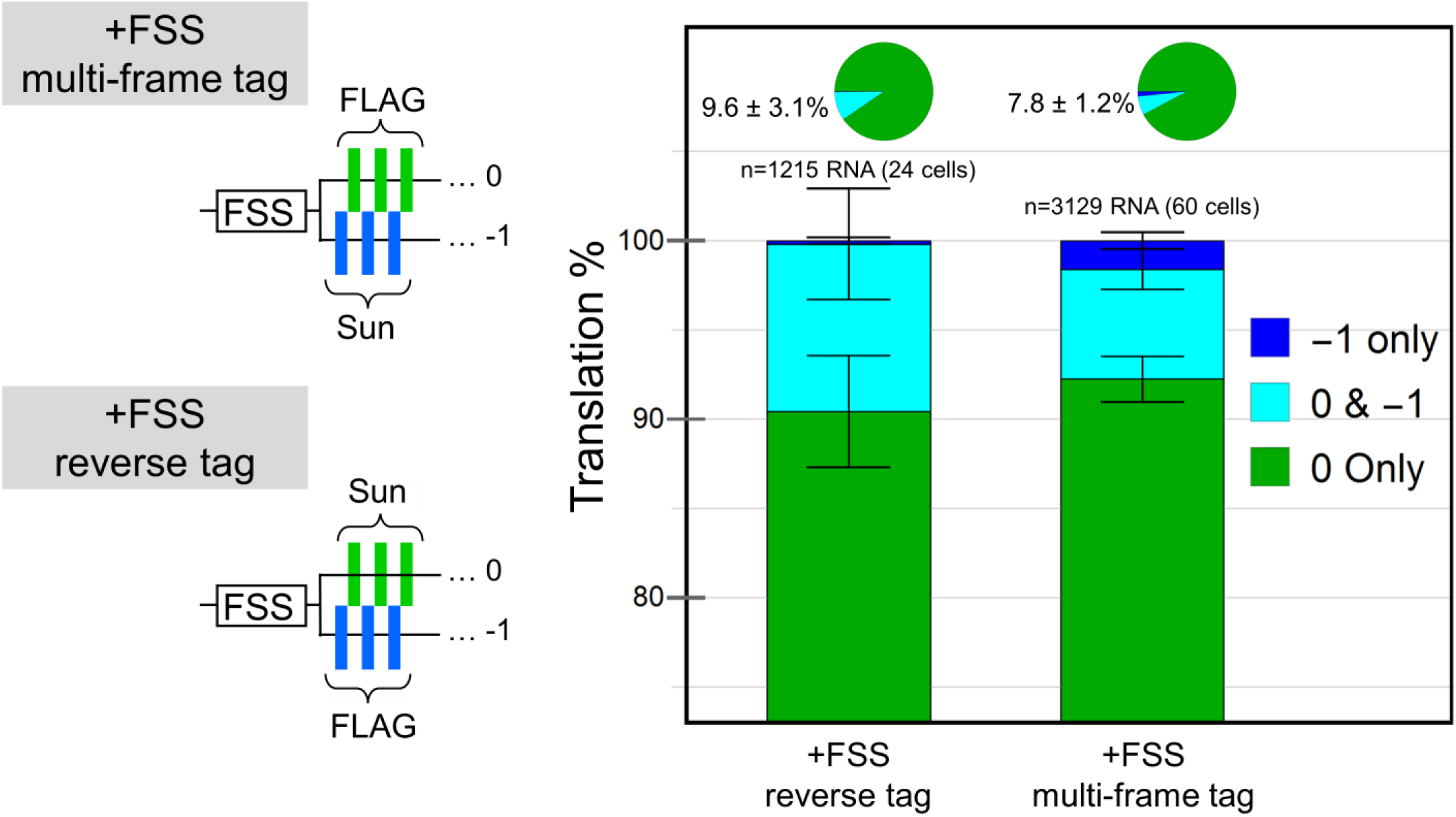
Epitope order in tags has little impact on percentage of frameshifting RNA. The percentage of RNA translating the −1 frame only (dark blue), the 0 and −1 frames (cyan), and the 0 frame only (green) for the +FSS multi-frame tag and the +FSS reverse tag (in which FLAG and SunTag epitopes are reversed). Pie charts show the percentage of translating species per cell. Error bars represent S.E.M. among cells.

**Figure S4.**
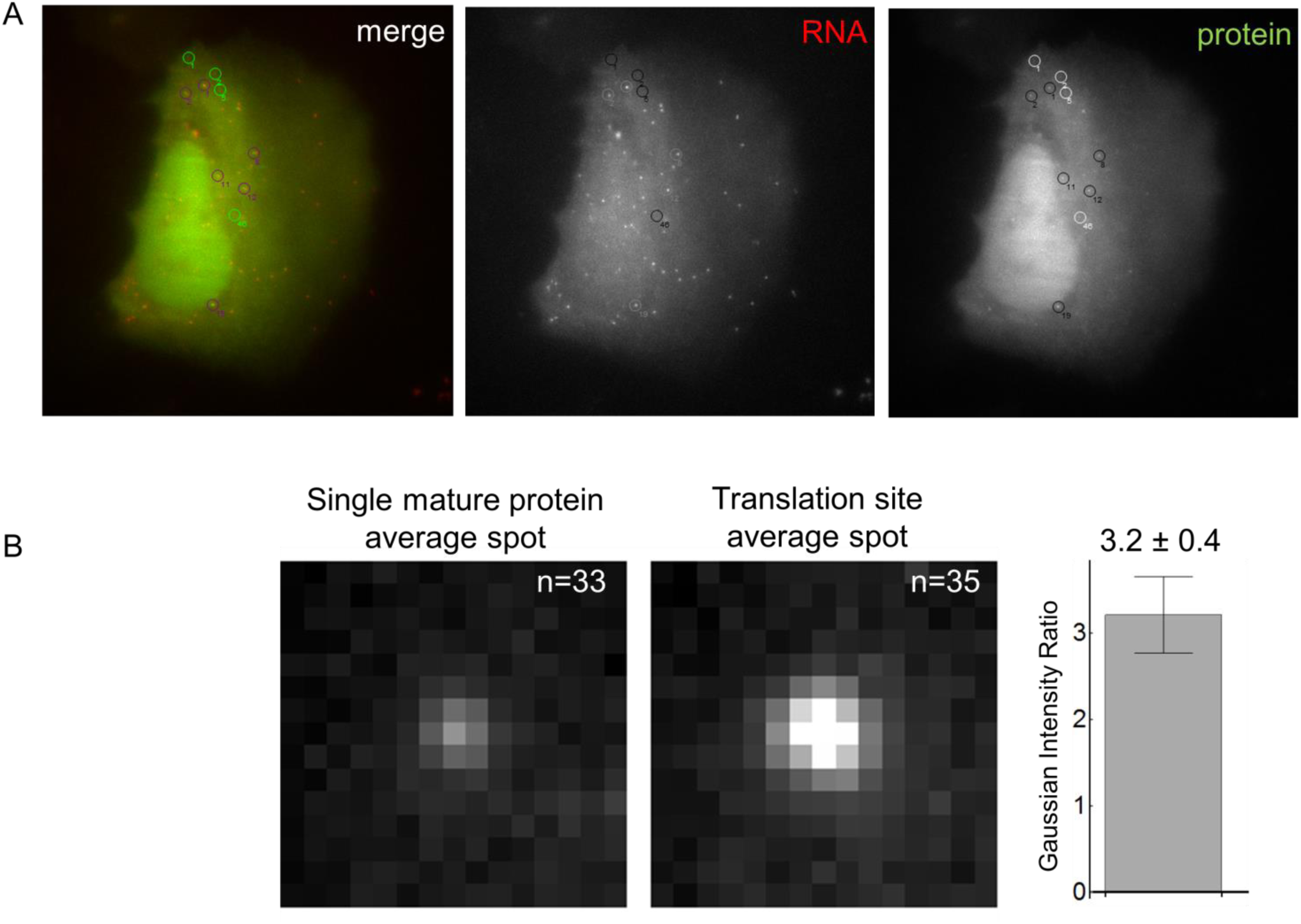
Calibrating translation site intensity to the number of ribosomes. (A) Example cell where single mature membrane proteins were detected (green circles), along with translation sites (co-localized RNA and protein; purple circles). (B) Average images of detected spots (33 mature proteins and 35 translation sites) were fit to a Gaussian to determine their intensity ratio, shown in the bar graph on the right. This ratio provides a calibration factor to convert translation site intensites to the number of riboeoms. The error bar represents the propagated 90% confidence interval from the fit.

**Figure S5.**
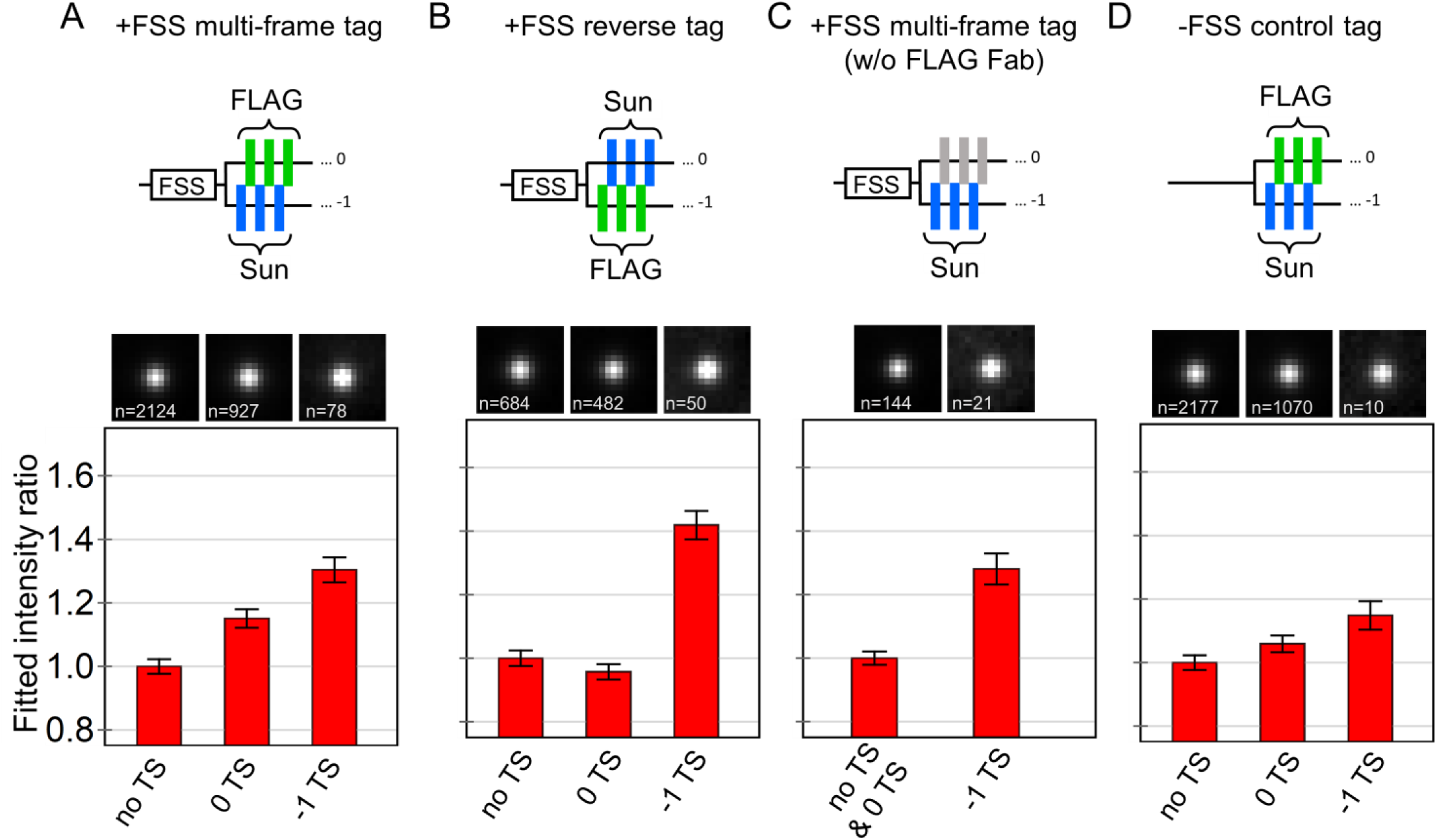
Frameshifting sites have brighter RNA signals. (A-E) Average RNA signals for non-translating RNA (no TS), 0 only translation sites (0 TS), and −1 frameshifting translation sites (−1 TS). The number of RNA used to generate each average image is shown. The bar graphs below show the Gaussian fit intensity (normalized to non-translating RNA, i.e. 0 TS sites). Experiments were done using (A) the +FSS multi-frame tag, (B) the +FSS reverse tag, (C) the +FSS multi-frame tag imaged without anti-FLAG Fab, and (D) the -FSS control tag. Error bars represent the fitted 90% confidence interval. Aside from the -FSS control, frameshifting sites (−1 TS) have significantly brighter RNA signals compared to canonical translation sites (0 TS). Note the frameshifting observed in the -FSS control represents less than 1% of translating RNA and can be considered background frameshifting.

**Figure S6.**
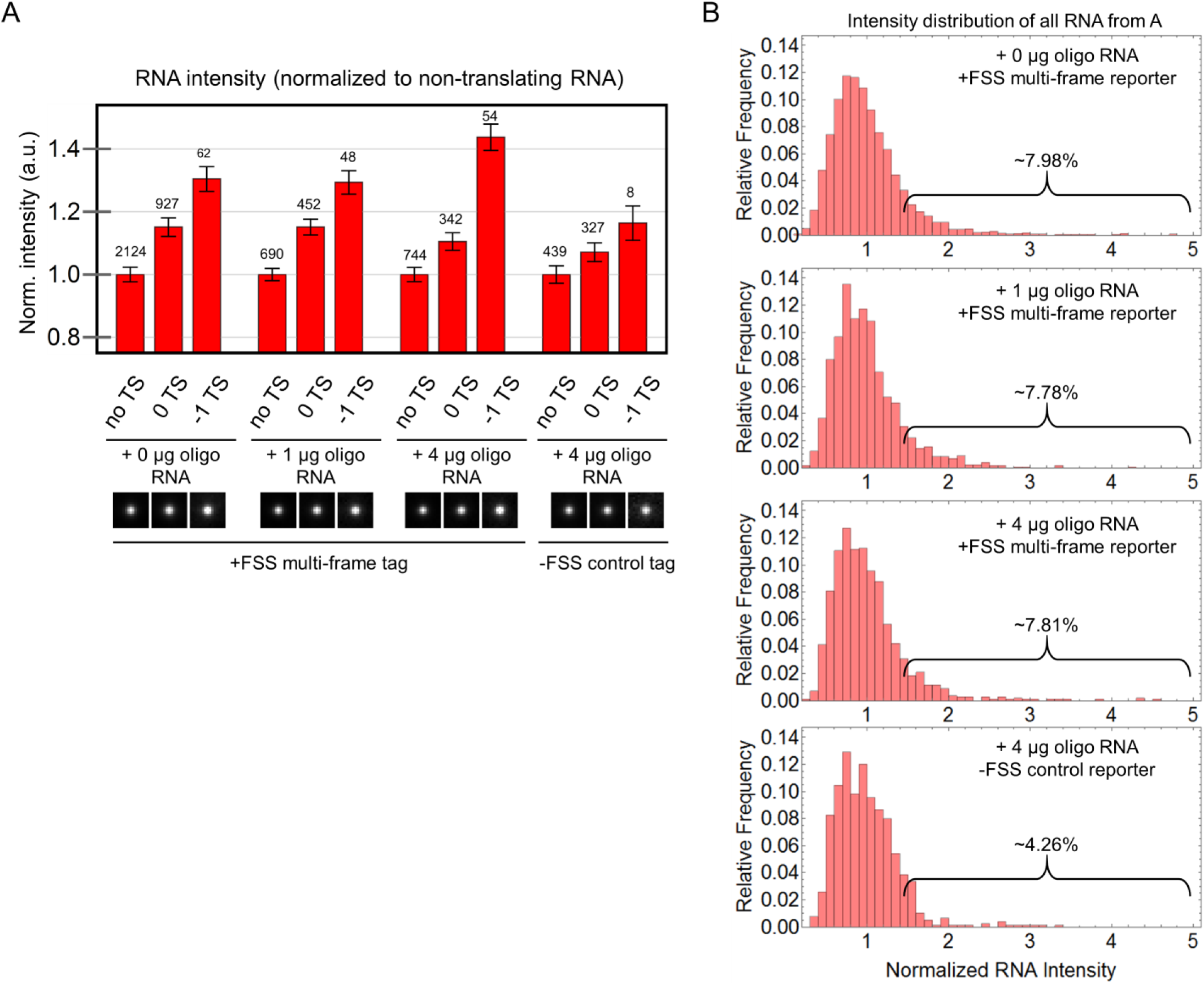
Upon oligo RNA co-transfection, stimulated frameshifting still occurs at bright RNA sites, even though the fraction of bright RNA sites remains unchanged. (A) Average RNA signals from non-translating RNA (no TS), 0-frame only translation sites (0 TS), and frameshifting translation sites (−1 TS) when different concentrations of oligo RNA (0, 1, and 4 mg) encoding the frameshift sequence were co-transfected into cells with the +FSS multi-frame reporter or the control -FSS reporter. The bar graph above shows the Gaussian fit intensity (normalized to non-translating RNA, i.e. 0 TS sites). (B) The distributions of RNA signals from the four experiments normalized to their mean value. In all +FSS experiments, the fraction of bright RNA (defined here as having an intensity greater than or equal to 1.6 times the mean) remained constant. The -FSS experiment had fewer bright RNA, suggesting the FSS sequence facilitates incorporation into multi-RNA sites.

**Figure S7.**
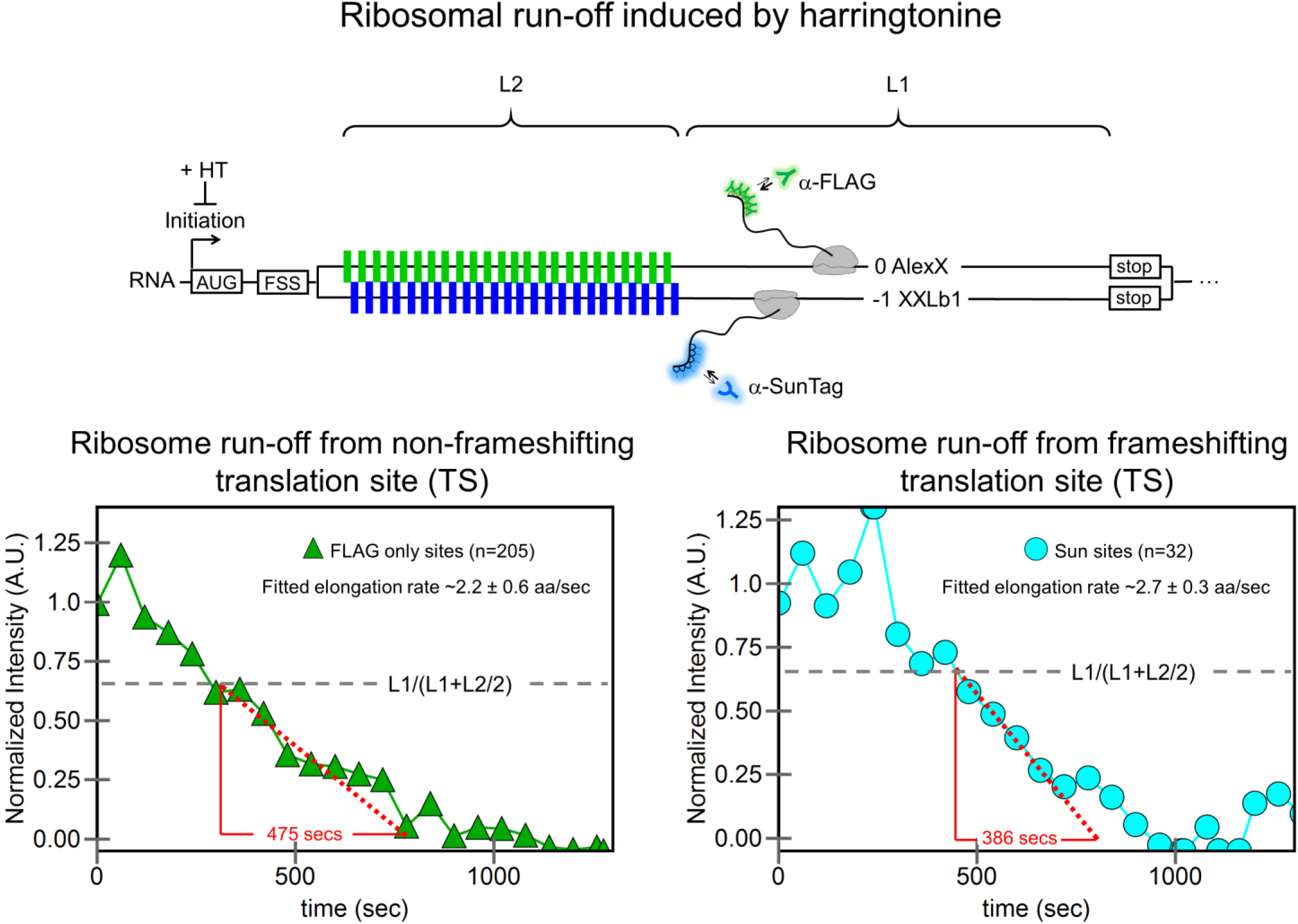
Fits to linear portion of ribosomal run-offs provide elongation rate estimates. The total intensity of detected non-frameshifting and frameshifting translation sites decays with time after addition (at t = 0 sec) of the translational initiation inhibitor harringtonine. Data is taken from Fig. 4A. If L2 is the length of the tagged portion of the open reading frame and L1 is the length of the non-tagged portion, then the linear (post-tag) portion of the decay begins from approximately L1/(L1+L2/2). This portion of the curve provides an estimate of the elongation rate during the part of the run-off where no new epitopes are being translated. Therefore, the run-off fit is independent of the frameshifting kinetics that occur upstream of the epitopes. Curves are normalized to their initial values.

**Figure S8.**
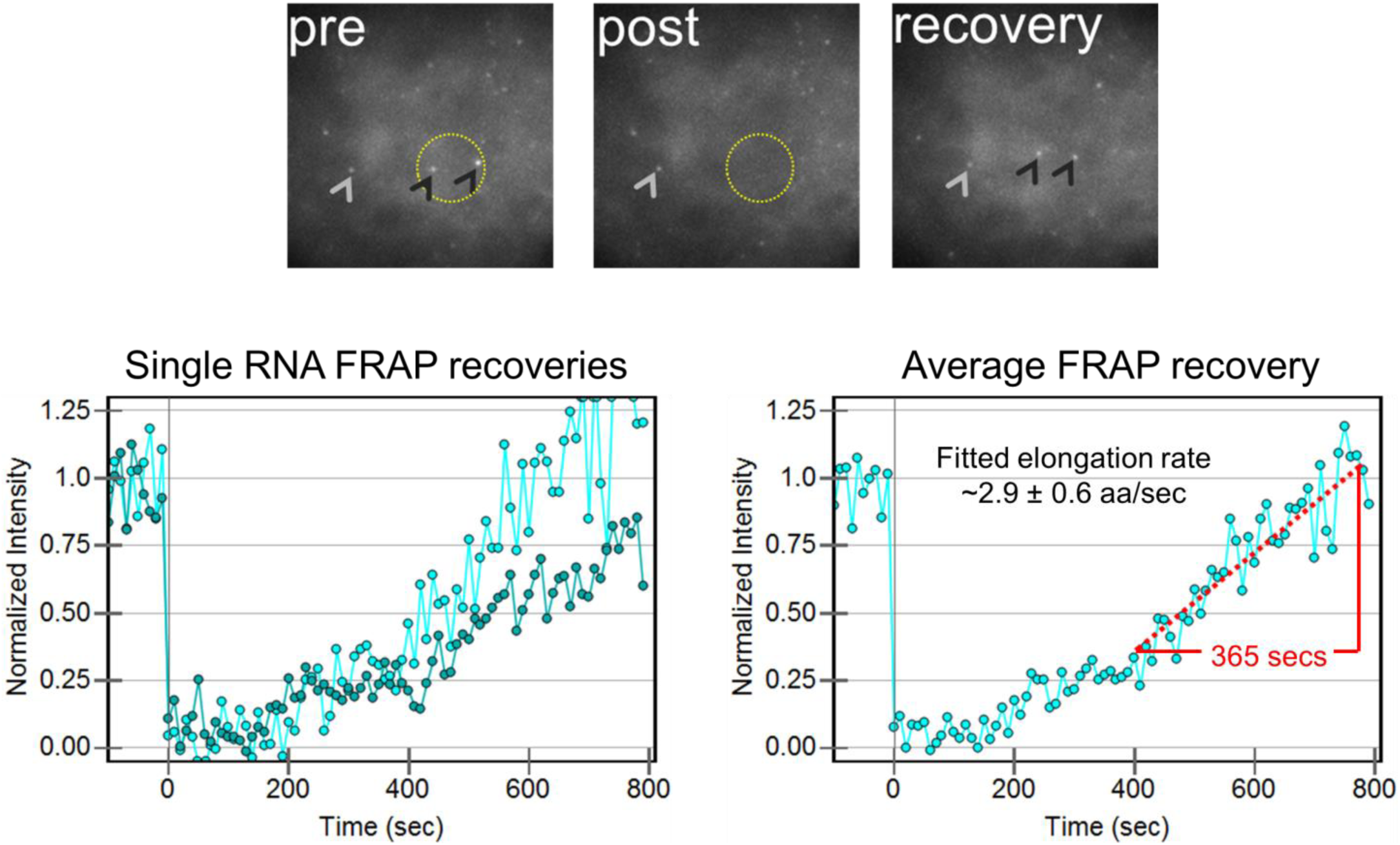
Fluroescence recovery after photobleach of frameshifting sites. (A) Fluorescence recovery after photobleaching (FRAP) experiments were performed at frameshifting translation sites (yellow circle marks the photobleach spot). The fluorescence recovery of the 0-frame signal within these sites was quantified as a function of time, with sample pre, post, and recovery frames shown above. (B) The average FRAP recovery time can be fit to estimate the elongation rate. This rate is similar to what was measured with harringtonine in Fig. S7.

**Figure S9.**
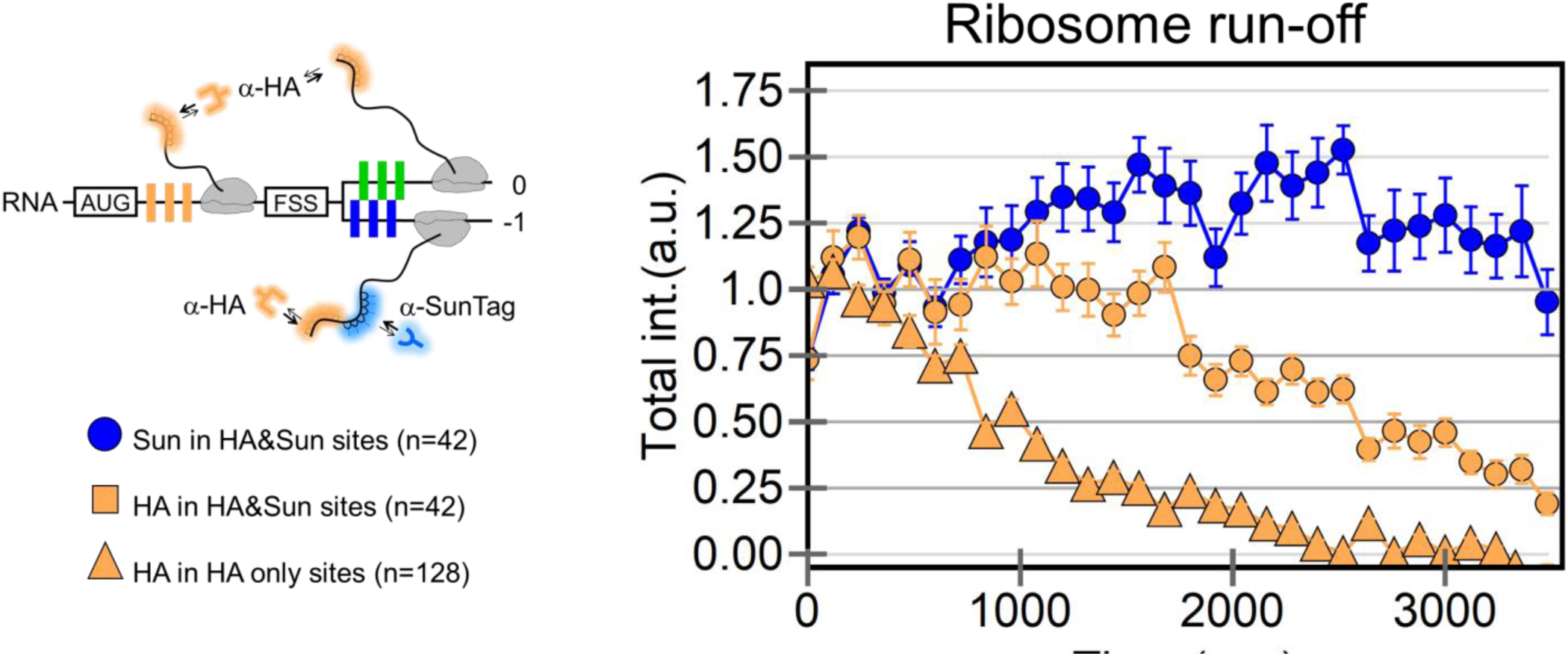
Ribosome run-off from the HA multi-frame tag. Ribosome run-off curve showing Sun signal in HA&Sun sites from the experiment performed in Figure 4B. The Sun signal comes from frameshifted ribosomes that have run past the frameshift sequence (FSS). Error bars represent S.E.M.

**Figure S10.**
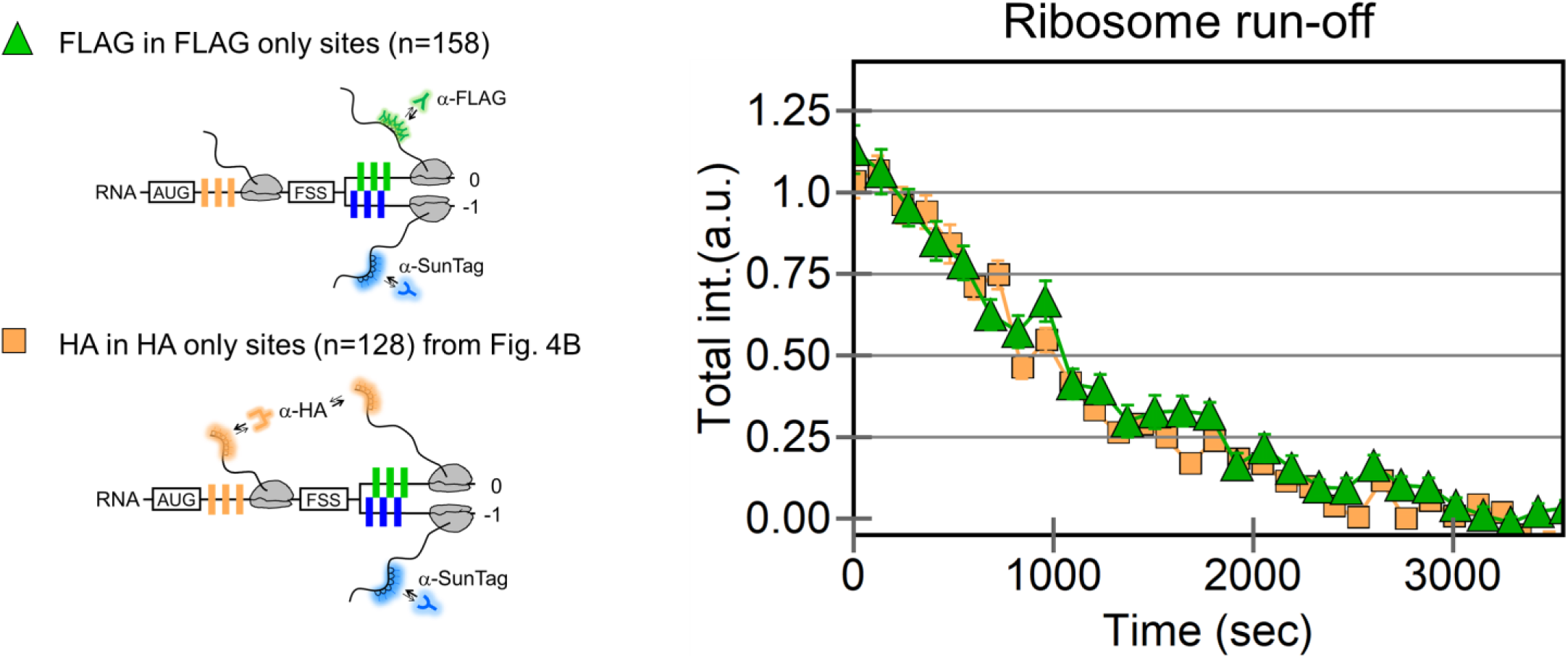
Ribosome run-off from FLAG epitopes in HA multi-frame tag. For completeness, an additional experiment was performed to generate the ribosome run-off curve from FLAG epitopes in the HA multi-frame tag. Because we can only image two epitopes at the same time (since RNA is imaged in a third color), we examined FLAG and Sun epitopes (Sun epitopes were required to distinguish frameshifting and non-frameshifting sites). The FLAG run-off from this experiment is on the right (green triangles). This curve is compared to the run-off curve of HA in HA only sites (orange squares) from Figure 4B. Both experiments represent non-frameshifting sites (no Sun signal detected). Error bars represent S.E.M.

**Figure S11.**
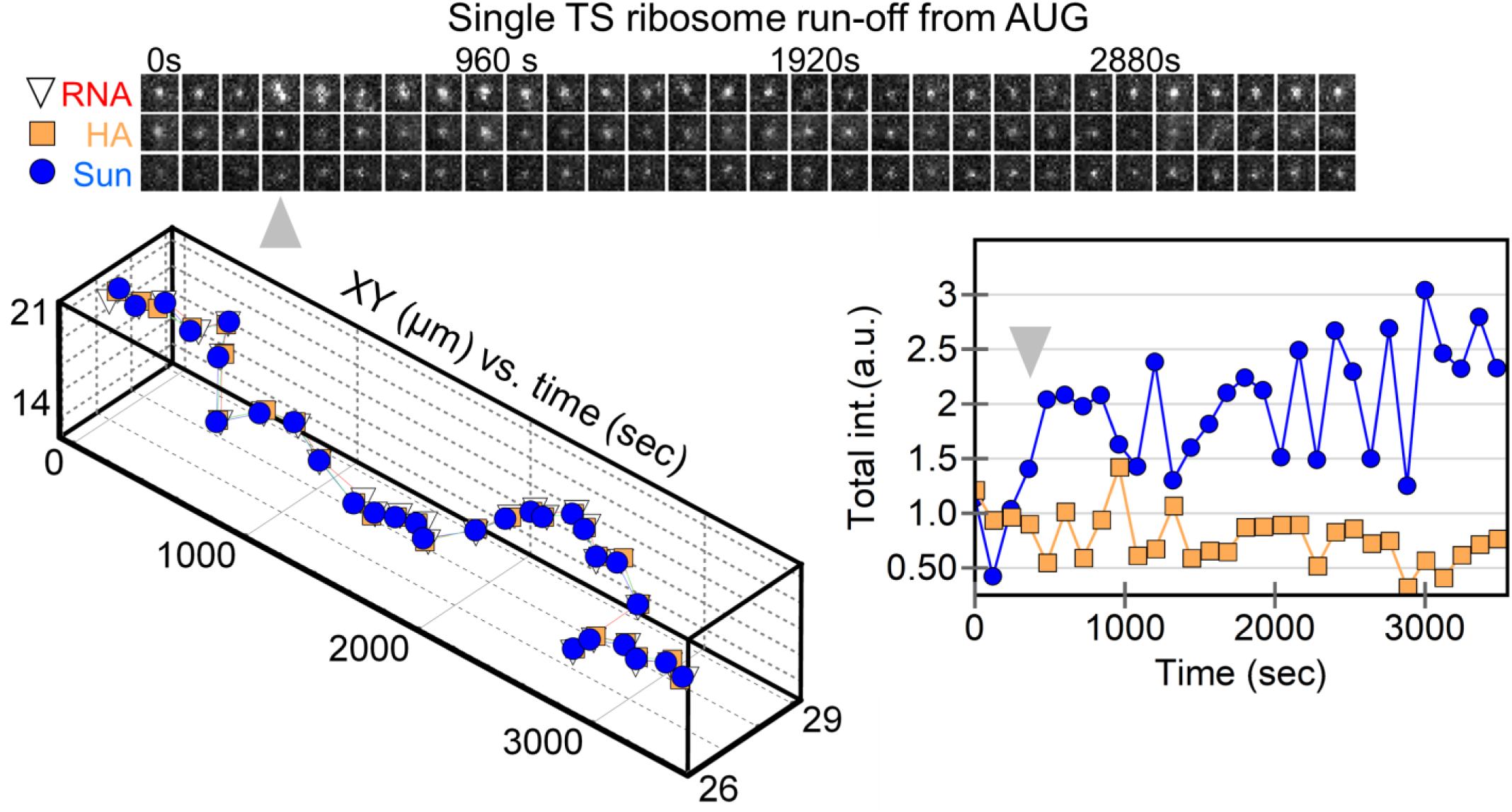
Track of a stimulated frameshifting burst at a single translation site. A single frameshifting translation site encoding the HA multi-frame tag was tracked after harringtonine addition. On the top, a montage of image trims shows the detected RNA-, HA-, and Sun shown signals through time. Below, the positions of the detected signals within the site are plotted through time. On the right, the normalized total intensity of the HA Fab signal (marking all ribosomes) and the SunTag scFv signal (marking frameshifting ribosomes) is plotted through time. At about the fourth timepoint, another non-translating RNA interacts with the frameshifting RNA, leading to a burst of frameshifting signal (Sun). Gray arrows signify a burst of frameshifting, coinciding with multi-RNA interactions.

**Figure S12.**
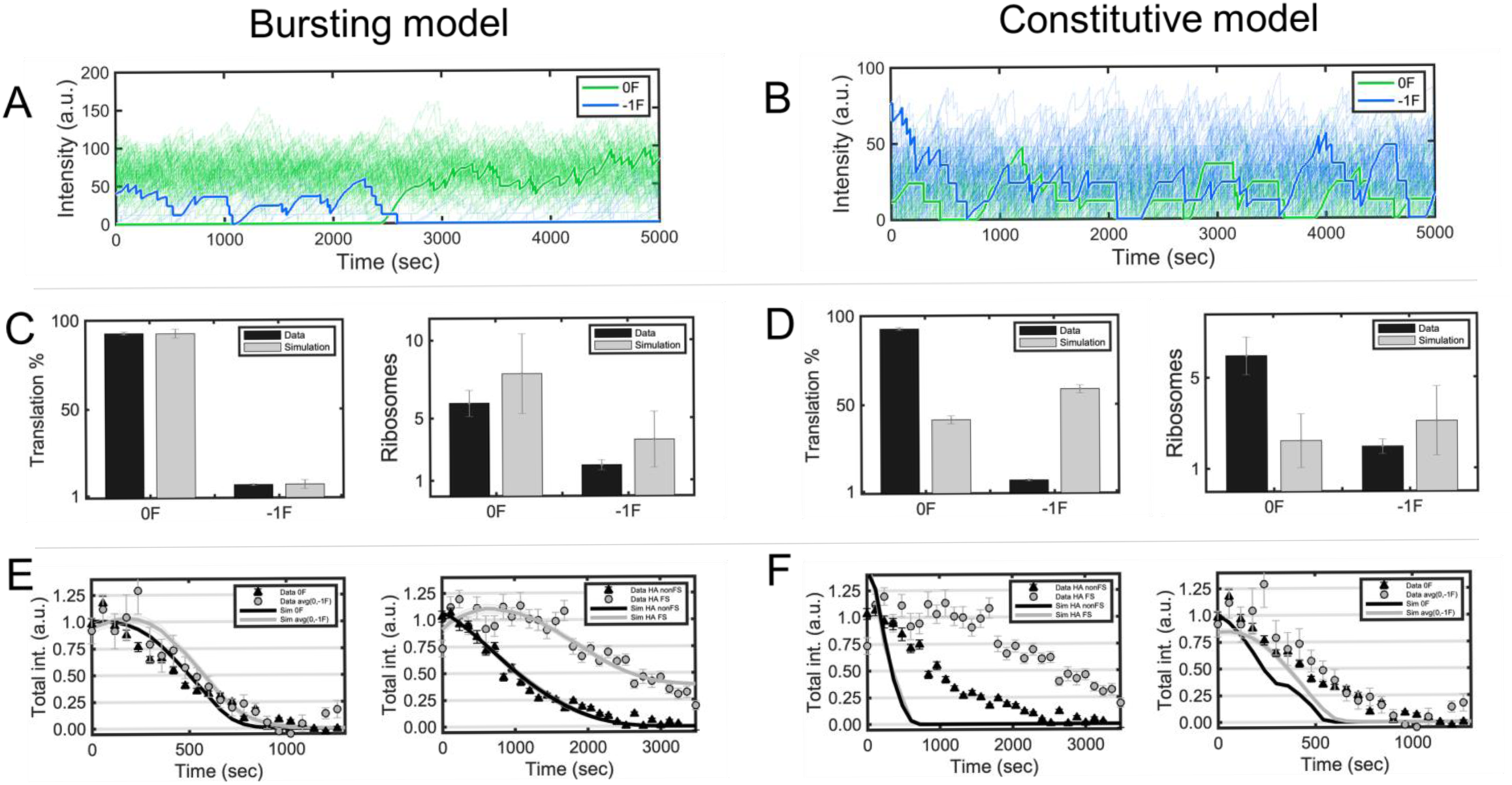
Comparison between the bursting and constitutive models. Model dynamics for the bursting (A,C,E) and constitutive model (B,D,F). Simulations were performed using the best parameter values obtained from the optimization process. (A,B) Simulated time courses representing single molecule fluctuation dynamics from 100 translation sites (a sample trace is shown in bold). Green and blue lines represent the translation of FLAG (0 frame) and Sun (−1 frame) epitopes in the +FSS multi-frame tag, respectively. (C,D) On the left, bar graphs showing the experimental (black) and simulated (gray) percentages of 0 and −1 frame translation. On the right, the number of ribosomes translating either the 0 or −1 frame. (E,F) On the left, simulated run-off (solid lines) from the frameshift sequence (FSS) of all ribosomes in non-frameshifting (black) and frameshifting sites (gray), plotted with data in Fig. 4A (black triangles and gray circles). On the right, simulated run-off (solid lines) from the start site (AUG) of all ribosomes in non-frameshifting (black) and frameshifting sites (gray), plotted with data in Fig. 4B (black triangles and gray circles). Error bars represent the standard error of the mean (S.E.M.). Details can be found in the Supplementary Methods.

**Figure S13.**
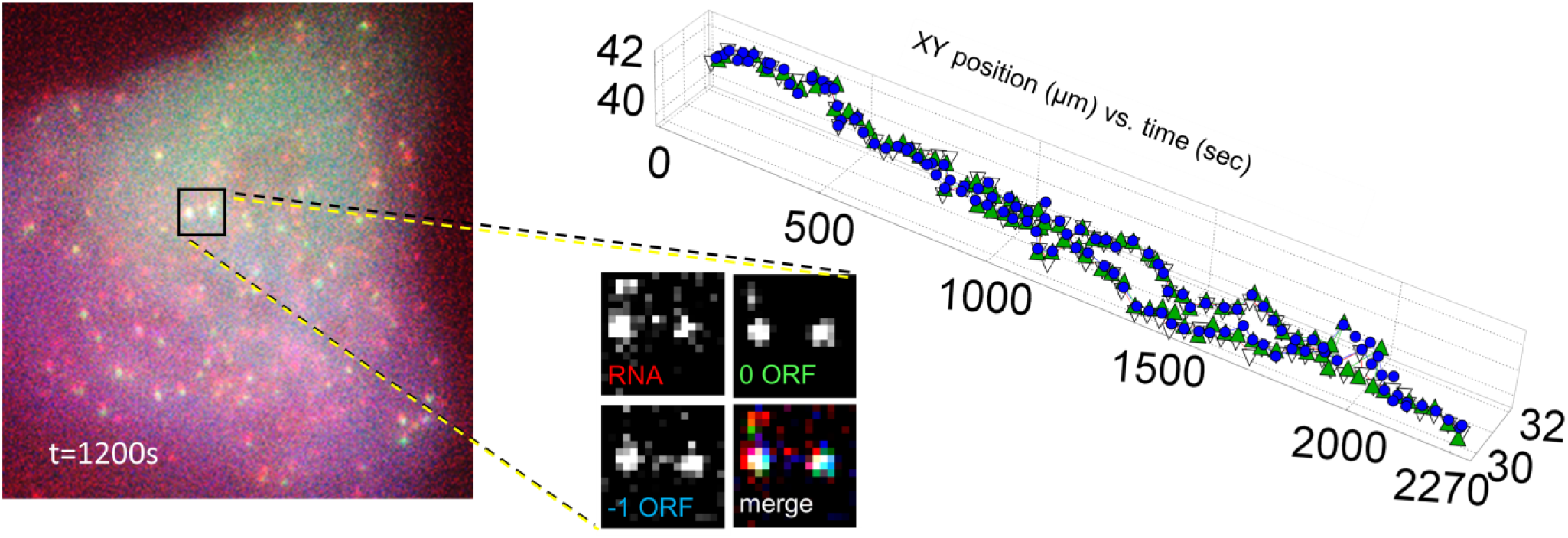
Single frameshifting RNA track persisting for >35 minutes. A sample frameshifting translation site encoding the +FSS 2x multi-frame tag was tracked for >35 minutes. Left shows the cell at the 1200 second time point. The zoom shows the channels separated to highlight two frameshifted RNA that split from a single translation site. On the right, the position of the spot is plotted through time.

**Figure S14.**
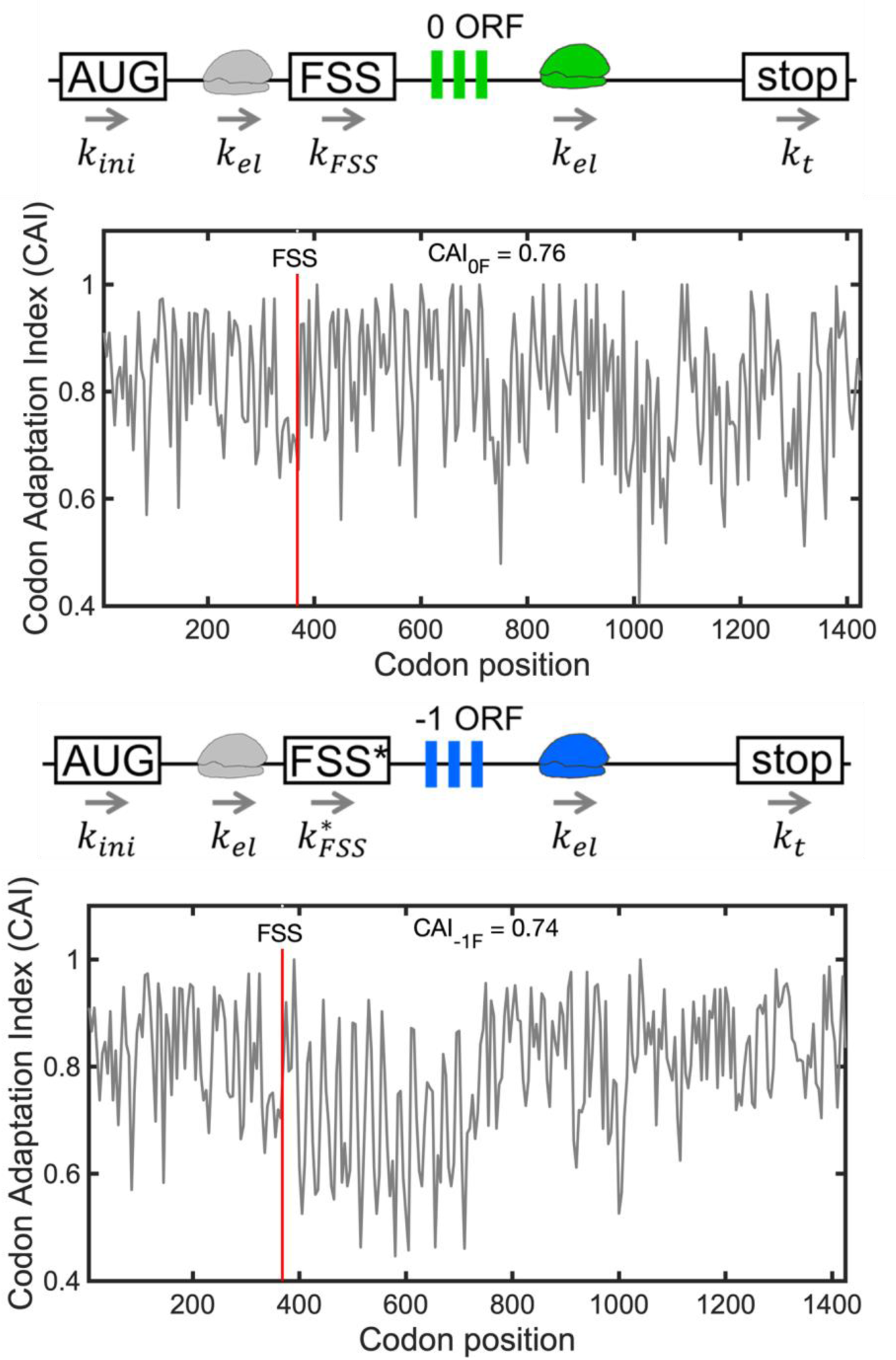
Codon usage plays a minor role in traffic jam formation. (A) The codon adaptation index (CAI) for the 0 frame (top) and −1 frame (bottom) of the +FSS multi-frame tag. The y-axis shows the CAI calculated using the codon frequency in the human genome. The x-axis shows the length of the genes, in codons. In the plots, rare codons have low CAIs and common codons have high CAIs. The vertical red line represents the location of the frameshift sequence (FSS). Similar CAIs are obtained for the sequences in the 0 frame (CAI_0F_ = 0.76) and the −1 frame (CAI_-1F_ = 0.74).

**Figure S15.**
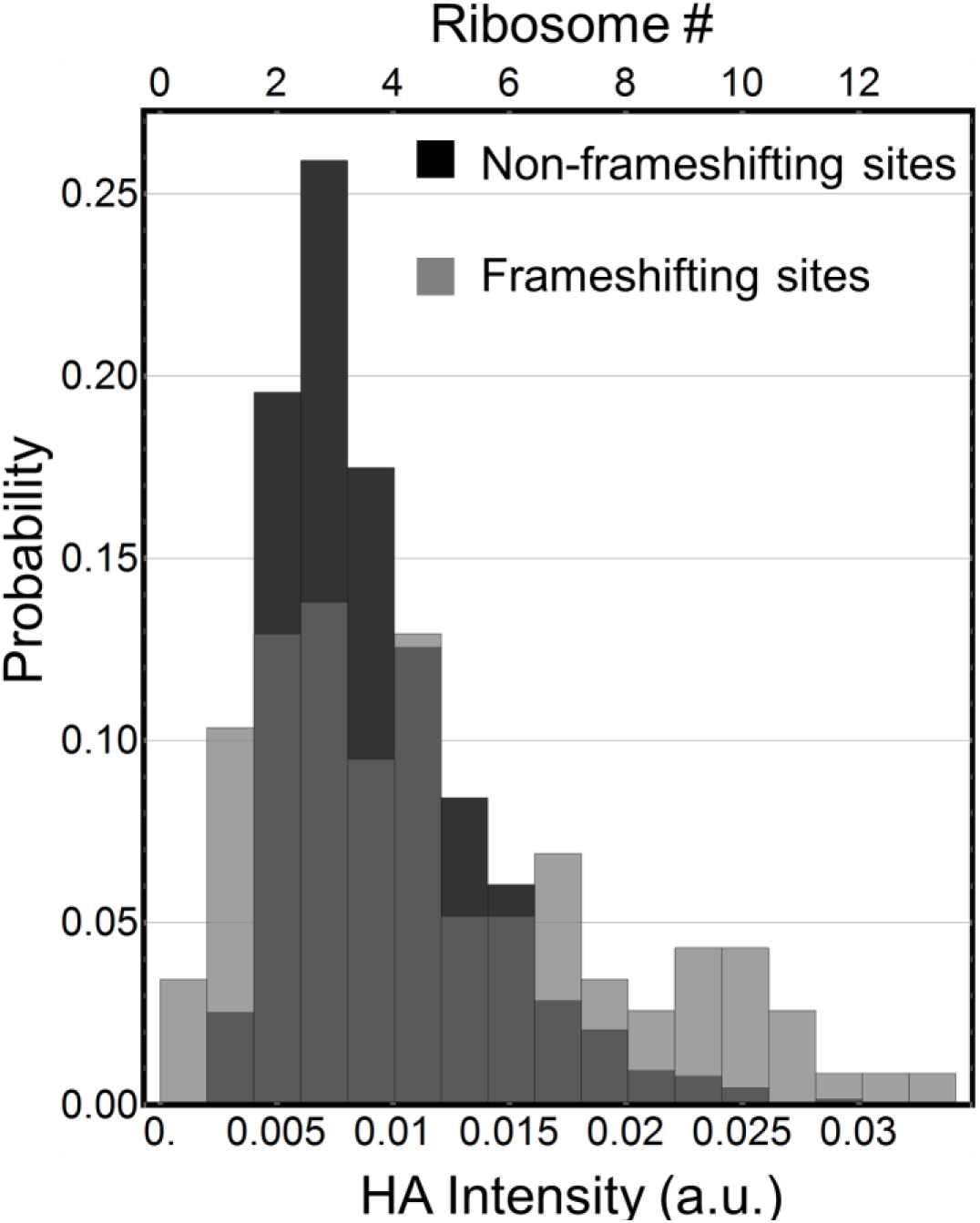
HA signal intensity distributions of non-frameshifting and frameshifting sites. Intensity distribution of non-frameshifted HA signals (black) versus frameshifted HA signals (gray) produced from the HA multi-frame tag.

